# *Staphylococcus aureus* uses the bacilliredoxin (BrxAB)/ bacillithiol disulfide reductase (YpdA) redox pathway to defend against oxidative stress under infections

**DOI:** 10.1101/624676

**Authors:** Nico Linzner, Vu Van Loi, Verena Nadin Fritsch, Quach Ngoc Tung, Saskia Stenzel, Markus Wirtz, Rüdiger Hell, Chris Hamilton, Karsten Tedin, Marcus Fulde, Haike Antelmann

**Author notes:** **Corresponding author:** Haike Antelmann, Institute for Biology-Microbiology, Freie Universität Berlin, Königin-Luise-Strasse 12-16, D-14195 Berlin, Germany.

## Abstract

*Staphylococcus aureus* is a major human pathogen and has to cope with reactive oxygen and chlorine species (ROS, RCS) during infections. The low molecular weight thiol bacillithiol (BSH) is an important defense mechanism of *S. aureus* for detoxification of ROS and HOCl stress to maintain the reduced state of the cytoplasm. Under HOCl stress, BSH forms mixed disulfides with proteins, termed as *S*-bacillithiolations, which are reduced by bacilliredoxins (BrxA and BrxB). The NADPH-dependent flavin disulfide reductase YpdA is phylogenetically associated with the BSH synthesis and BrxA/B enzymes and was proposed to function as BSSB reductase. Here, we investigated the role of the bacilliredoxin BrxAB/BSH/YpdA pathway in *S. aureus* COL under oxidative stress and macrophage infection conditions *in vivo* and in biochemical assays *in vitro*. Using HPLC thiol metabolomics, a strongly enhanced BSSB level and a decreased BSH/BSSB ratio were measured in the *S. aureus* COL *ypdA* deletion mutant under control and NaOCl stress. Monitoring the BSH redox potential (*E*_BSH_) using the Brx-roGFP2 biosensor revealed that YpdA is required for regeneration of the reduced *E*_BSH_ upon recovery from oxidative stress. In addition, the *ypdA* mutant was impaired in H_2_O_2_ detoxification as measured with the novel H_2_O_2_-specific Tpx-roGFP2 biosensor. Phenotype analyses further showed that BrxA and YpdA are required for survival under NaOCl and H_2_O_2_ stress *in vitro* and inside murine J-774A.1 macrophages in infection assays *in vivo*. Finally, NADPH-coupled electron transfer assays provide evidence for the function of YpdA in BSSB reduction, which depends on the conserved Cys14 residue. YpdA acts together with BrxA and BSH in de-bacillithiolation of *S*-bacilithiolated GapDH. In conclusion, our results point to a major role of the BrxA/BSH/YpdA pathway in BSH redox homeostasis in *S. aureus* during recovery from oxidative stress and under infections.

## INTRODUCTION

*Staphylococcus aureus* is an important human pathogen, which can cause many diseases, ranging from local soft-tissue and wound infections to life-threatening systemic and chronic infections, such as endocarditis, septicaemia, bacteraemia, pneumonia or osteomyelitis (Archer, 1998;Lowy, 1998;Boucher and Corey, 2008). Due to the prevalence of methicillin-resistant *S. aureus* isolates, which are often resistant to multiple antibiotics, treatment options are limited to combat *S. aureus* infections (Livermore, 2000). Therefore, the “European Center of Disease Prevention and Control” has classified *S. aureus* as one out of six ESKAPE pathogens which are the leading causes of nosocomial infections worldwide (Pendleton et al., 2013). During infections, activated macrophages and neutrophils produce reactive oxygen and chlorine species (ROS, RCS) in large quantities, including H_2_O_2_ and hypochlorous acid (HOCl) with the aim to kill invading pathogens (Winterbourn and Kettle, 2013;Hillion and Antelmann, 2015;Beavers and Skaar, 2016;Winterbourn et al., 2016).

*S. aureus* has several defense mechanisms against ROS and HOCl, such as the MarR/SarA family redox regulators SarZ and MgrA, the HOCl-specific Rrf2-family regulator HypR and the peroxide-specific PerR repressor (Horsburgh et al., 2001;Chen et al., 2006;Chen et al., 2009;Chen et al., 2011;Hillion and Antelmann, 2015;Ji et al., 2015;Chandrangsu et al., 2018;Loi et al., 2018b). In addition, low molecular weight (LMW) thiols play important roles for survival and pathogenicity in bacterial pathogens (Loi et al., 2015;Tung et al., 2018). Gram-negative bacteria produce glutathione (GSH) as major LMW thiol, which is absent in most Gram-positive bacteria (Fahey, 2013). Instead, many firmicutes utilize bacillithiol (BSH) as alternative LMW thiol, which is essential for virulence of *S. aureus* in macrophage infection assays (Newton et al., 2012;Pöther et al., 2013;Posada et al., 2014;Chandrangsu et al., 2018). A recent study identified a BSH derivative with an N-methylated cysteine as N-methyl-BSH in anaerobic phototrophic *Chlorobiaceae*, suggesting that BSH derivatives are more widely distributed and not restricted to Gram-positive firmicutes (Hiras et al., 2018). In *S. aureus* and *Bacillus subtilis*, BSH was characterized as cofactor of thiol-*S*-transferases (e.g. FosB), glyoxalases, peroxidases and other redox enzymes that are involved in detoxification of ROS, HOCl, methylglyoxal, toxins and antibiotics (Chandrangsu et al., 2018). In addition, BSH participates in post-translational thiol-modifications under HOCl stress by formation of BSH mixed protein disulfides, termed as protein *S*-bacillithiolations (Chi et al., 2011;Chi et al., 2013;Imber et al., 2018a;Imber et al., 2018c).

*S*-bacillithiolated proteins are widespread and conserved across firmicutes and were discovered in many *Bacillus* and *Staphylococcus* species (Chi et al., 2013). Protein *S*-bacillithiolation functions in thiol-protection and redox regulation of redox-sensing regulators, metabolic enzymes and antioxidant enzymes (Chi et al., 2011;Chi et al., 2013;Loi et al., 2015;Imber et al., 2018a;Imber et al., 2018b;Imber et al., 2018c). In *S. aureus*, the glycolytic glyceraldehyde-3-phosphate dehydrogenase (GapDH) and the aldehyde dehydrogenase AldA were identified as most abundant proteins that are *S*-bacillithiolated under HOCl stress (Imber et al., 2018a;Imber et al., 2018b). Using OxICAT analyses, GapDH and AldA displayed also the highest oxidation increase of 29% in the *S. aureus* redox proteome upon HOCl stress. GapDH activity was inhibited by *S*-bacillithiolation of its active site Cys151 under H_2_O_2_ and NaOCl stress *in vitro*, which was faster compared to the irreversible inactivation by overoxidation to Cys sulfonic acids (Imber et al., 2018a). These experiments revealed that *S*-bacillithiolation efficiently protects the GapDH active site against overoxidation (Loi et al., 2015;Imber et al., 2018c).

In *B. subtilis*, the methionine synthase MetE was *S*-bacillithiolated at its Zn binding active site Cys730 under HOCl stress leading to its inactivation and methionine auxotrophy (Chi et al., 2011). In addition, the MarR-family organic hydroperoxide repressor OhrR is *S*-bacillithiolated at its redox-sensing single Cys under cumene hydroperoxide and HOCl stress (Fuangthong et al., 2001;Lee et al., 2007;Chi et al., 2011). *S*-bacillithiolation leads to inhibition of OhrR repressor activity and derepression of the OhrA peroxiredoxin for detoxification of organic hydroperoxides that could originate from fatty acid oxidation (Dubbs and Mongkolsuk, 2007). The OhrR-homologs SarZ and MgrA of *S. aureus* were also shown to be redox-regulated by *S*-thiolation using benzene thiol *in vitro* based on crystal structure analyses (Abomoelak et al., 2009;Poor et al., 2009;Chen et al., 2011;Sun et al., 2012;Hillion and Antelmann, 2015). In addition, OxICAT analyses revealed an >10% oxidation increase of MgrA and SarZ under HOCl stress in *S. aureus* indicating their possible redox-regulation by *S*-bacillithiolation *in vivo* (Imber et al., 2018a).

Reduction of *S*-bacillithiolated OhrR, MetE and GapDH proteins is catalyzed by the bacilliredoxins (BrxA/B) in *B. subtilis* and *S. aureus in vitro* (Gaballa et al., 2014;Chandrangsu et al., 2018). BrxA (YphP) and BrxB (YqiW) are paralogous thioredoxin-fold proteins of the UPF0403 family with an unusual CGC active site **(Fig. S1)** (Derewenda et al., 2009;Gaballa et al., 2014). Upon de-bacillithiolation, the BSH moiety is transferred to the Brx active site, resulting in BrxA-SSB formation (Fig. 1B). However, the Brx associated thiol-disulfide reductase involved in regeneration of Brx activity is not known. In GSH-producing bacteria, glutaredoxins (Grx) catalyze the reduction of *S*-glutathionylated proteins, which requires GSH for regeneration of Grx, resulting in glutathione disulfide (GSSG) formation (Lillig et al., 2008;Allen and Mieyal, 2012). The regeneration of GSH is catalyzed by the flavoenzyme glutathione disulfide reductase (Gor), which belongs to the pyridine nucleotide disulfide reductases and recycles GSSG on expense of NADPH (Argyrou and Blanchard, 2004).

**Fig. 1.**
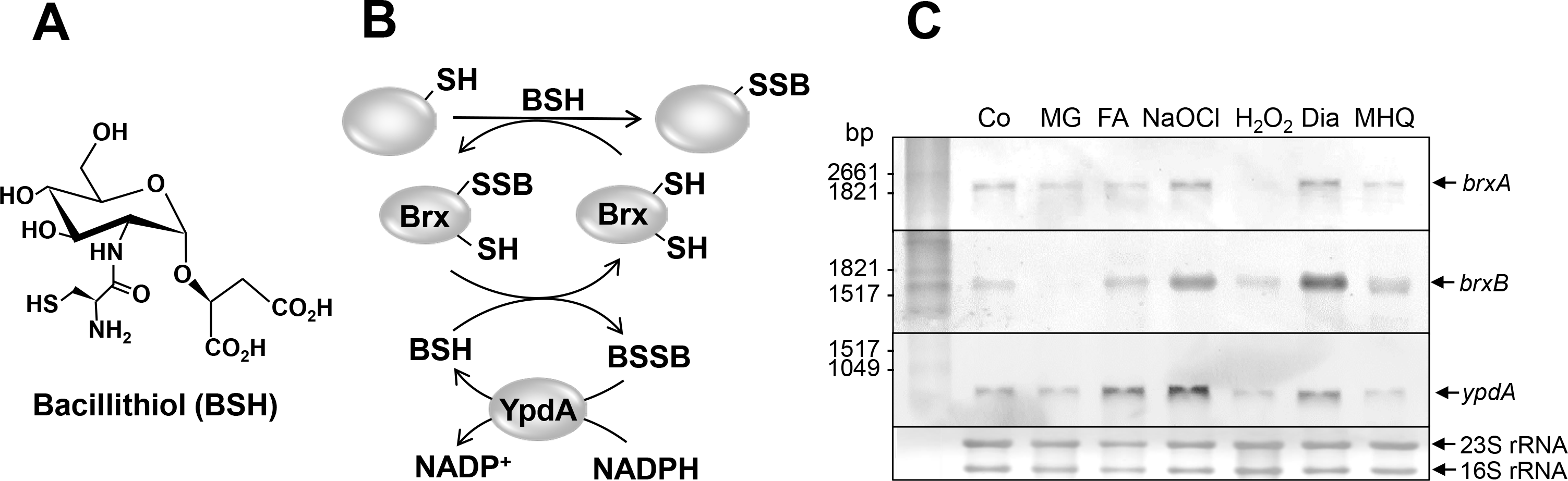
Structure of the LMW thiol bacillithiol (BSH), proposed mechanism of the BrxAB/BSH/YpdA de-bacillithiolation pathway and induction of the *brxA*,*brxB* and *ypdA* genes and operons in *S. aureus* COL. **(A)** Bacillithiol is the major LMW thiol in *S. aureus.* **(B)** Under HOCl stress, BSH forms mixed disulfides with proteins, termed as *S*-bacillithiolations (SSB). Bacilliredoxin (BrxA/B) are involved in reduction of *S*-bacillithiolated proteins, resulting in the transfer of BSH to the Brx active site (Brx-SSB). BSH functions in Brx-SSB reduction to restore Brx activity, leading to BSSB formation. The NADPH-dependent flavin disulfide reductase YpdA (SACOL1520) functions as BSSB reductase to regenerate BSH. **(C)** Northern blot analysis was used to analyze transcription of *brxA*, *brxB* and *ypdA* in *S. aureus* COL wild type before (Co) and 15 min after exposure to 0.5 mM methylglyoxal (MG), 0.75 mM formaldehyde (FA), 1 mM NaOCl, 10 mM H_2_O_2_, 2 mM diamide (Dia) and 45 μM methylhydroquinone (MHQ) stress at an OD_500_ of 0.5. The arrows point toward the transcript sizes of the *brxA*, *brxB* and *ypdA* specific genes and operons. The methylene blue stain is the RNA loading control indicating the 16S and 23S rRNAs.

An increased level of BSSB was previously measured under HOCl stress in *B. subtilis* using HPLC metabolomics (Chi et al., 2013). In *S. aureus*, HOCl and H_2_O_2_ stress resulted in an increased BSH redox potential as revealed using the Brx-roGFP2 biosensor, which is highly specific for BSSB (Loi et al., 2017). However, the identity of the BSSB reductase for regeneration of reduced BSH was unknown thus far. Phylogenetic profiling has suggested the flavoenzyme YpdA (SACOL1520) as putative NADPH-dependent BSSB reductase, which is encoded only in BSH producing bacteria **(Fig. S1)** (Gaballa et al., 2010). However, biochemical evidence for the function of YpdA as BSSB reductase is missing. Moreover, the physiological functions of the BrxA/B and YpdA enzymes in the oxidative stress response have not been demonstrated in *B. subtilis* and *S. aureus*. In this work, we used phenotype and biochemical analyses, HPLC metabolomics and redox biosensor measurements to study the physiological role of the Brx/BSH/YpdA redox pathway in *S. aureus* under oxidative stress and macrophage infection assays. Our data point to important roles of BrxA and YpdA in the oxidative stress defense for regeneration of reduced *E*_BSH_ and de-bacillithiolation upon recovery from oxidative stress. Biochemical assays further provide evidence for the function of YpdA as BSSB reductase, which acts in the BrxA/BSH/YpdA electron pathway in de-bacillithiolation of GapDH-SSB.

## MATERIALS AND METHODS

### Bacterial strains, growth and survival assays

Bacterial strains, plasmids and primers used in this study are listed in **Tables S1, S2 and S3**. For cloning and genetic manipulation, *Escherichia coli* was cultivated in LB medium. For stress experiments, *S. aureus* COL wild type and mutant strains were cultivated in LB, RPMI or Belitsky minimal medium and exposed to the different compounds during the exponential growth as described previously (Loi et al., 2017;Loi et al., 2018b). Sodium hypochlorite, methylglyoxal, diamide, methylhydroquinone, dithiothreitol, cumene hydroperoxide (80% w/v), H_2_O_2_ (35% w/v), and monobromobimane were purchased from Sigma Aldrich.

### Cloning, expression and purification of His-tagged Brx-roGFP2, Tpx-roGFP2, GapDH, BrxA, YpdA and YpdAC14A proteins in *E. coli*

Construction of plasmids pET11b-*brx-roGFP2* for expression of the Brx-roGFP2 biosensor was described previously (Loi et al., 2017). The pET11b-derived plasmids for overexpression of the His-tagged GapDH and BrxA (SACOL1321) proteins were generated previously (Imber et al., 2018a). The plasmid pET11b-*brx-roGFP2* was used as a template for construction of the Tpx-roGFP2 biosensor to replace *brx* by the *tpx* gene of *S. aureus*. The *tpx* gene (*SACOL1762*) was PCR-amplified from chromosomal DNA of *S. aureus* COL using primers pET-tpx-for-NheI and pET-tpx-rev-SpeI **(Table S3)**, digested with *Nhe*I and *Bam*HI and cloned into plasmid pET11b-*brx-roGFP2* to generate pET11b-*tpx-roGFP2*. To construct plasmids pET11b-*ypdA* or pET11b-*ypdAC14A*, the *ypdA* gene (*SACOL1520*) was PCR-amplified from chromosomal DNA of *S. aureus* COL with pET-ypdA-for-NdeI or pET-ypdAC14A-for-NdeI as forward primers and pET-ypdA-rev-BamHI as reverse primer **(Table S3)**, digested with *Nde*I and *Bam*HI and inserted into plasmid pET11b (Novagen). For expression of His-tagged proteins (GapDH, BrxA, YpdA, YpdAC14A, Tpx-roGFP2), *E. coli* BL21(DE3) *plys*S carrying plasmids pET11b-*gap*, pET11b-*brxA*, pET11b-*ypdA*, pET11b-*ypdAC14A* and pET11b-*tpx-roGFP2* was cultivated in 1 l LB medium until an OD_600_ of 0.8 followed by addition of 1 mM IPTG (isopropyl-β-D-thiogalactopyranoside) for 16 h at 25°C. His_6_-tagged GapDH, BrxA, YpdA, YpdAC14A and Tpx-roGFP2 proteins were purified using His Trap™ HP Ni-NTA columns (5 ml; GE Healthcare, Chalfont St Giles, UK) and the ÄKTA purifier liquid chromatography system (GE Healthcare) as described (Loi et al., 2018b).

### Construction of *S. aureus* COL *ypdA, brxAB* and *brxAB ypdA* deletion mutants and complemented mutant strains

*S. aureus* COL *ypdA* (*SACOL1520*), *brxA* (*SACOL1464*) and *brxB* (*SACOL1558*) single deletion mutants as well as the *brxA brxB* double and *brxA brxB ypdA* triple mutants were constructed using pMAD as described (Arnaud et al., 2004;Loi et al., 2018b). Briefly, the 500 bp up- and downstream regions of *ypdA*, *brxA* and *brxB* were amplified using gene-specific primers **(Table S3)**, fused by overlap extension PCR and ligated into the *Bgl*II and *Sal*I sites of plasmid pMAD. The pMAD constructs were electroporated into *S. aureus* RN4220 and further transduced into *S. aureus* COL using phage 81 (Rosenblum and Tyrone, 1964). The clean deletions of *ypdA*, *brxA* or *brxB* were selected after plasmid excision as described (Loi et al., 2018b). The *brxAB* and *brxAB ypdA* double and triple mutants were obtained by transduction and excision of pMAD-Δ*brxB* into the *brxA* mutant, leading to the *brxAB* deletion and of plasmid pMAD-Δ*ypdA* into the *brxAB* mutant, resulting in the *brxAB ypdA* knockout. For construction of *ypdA, brxA* and *brxB* complemented strains, the xylose-inducible ectopic *E. coli/ S. aureus* shuttle vector pRB473 was applied (Brückner et al., 1993). Primers pRB-ypdA, pRB-brxA and pRB-brxB (**Table S3**) were used for amplification of the genes, which were cloned into pRB473 after digestion with *Bam*HI and *Kpn*I to generate plasmids pRB473-*ypdA*, pRB473-*brxA* and pRB473-*brxB*, respectively. The pRB473 constructs were transduced into the *ypdA* and *brxAB* deletion mutants as described (Loi et al., 2017).

### Construction of Tpx-roGFP2 and Brx-roGFP2 biosensor fusions in *S. aureus* COL

*tpx-roGFP2* fusion was amplified from plasmid pET11b-*tpx-roGFP2* with primers pRB-tpx-roGFP2-for-BamHI and pRB-tpx-roGFP2-rev-SacI and digested with *Bam*HI and *Sac*I **(Table S3).** The PCR product was cloned into pRB473 generating plasmid pRB473-*tpx-roGFP2*. The biosensor plasmids pRB473-*tpx-roGFP2* and pRB473-*brx-roGFP2* were electroporated into *S. aureus* RN4220 and further transferred to the *S. aureus* COL *ypdA*, *brxAB* and *brxAB ypdA* mutants by phage transduction as described (Loi et al., 2017).

### Northern blot experiments

Northern blot analyses were performed using RNA isolated from *S. aureus* COL before and 15 min after exposure to 0.5 mM methylglyoxal, 0.75 mM formaldehyde, 1 mM NaOCl, 10 mM H_2_O_2_, 2 mM diamide, and 45 μM methylhydroquinone as described (Wetzstein et al., 1992). Hybridizations were conducted using digoxigenin-labeled antisense RNA probes for *ypdA, brxA* and *brxB* that were synthesized *in vitro* using T7 RNA polymerase and primers ypdA-NB-for/rev, brxA-NB-for/rev or brxB-NB-for/rev **(Table S3)** as in previous studies (Tam le et al., 2006).

### HPLC thiol metabolomics for quantification of LMW thiols and disulfides

For preparation of thiol metabolomics samples, *S. aureus* COL WT, *ypdA* and *brxAB* mutants as well as the *ypdA* complemented strains were grown in RPMI medium to an OD_500_ of 0.9 and exposed to 2 mM NaOCl stress for 30 min. The intracellular amounts of reduced and oxidized LMW thiols and disulfides (BSH, BSSB, cysteine and cystine) were extracted from the *S. aureus* cells, labelled with monobromobimane and measured by HPLC thiol metabolomics as described (Chi et al., 2013).

### Western blot analysis

*S. aureus* strains were grown in LB until an OD_540_ of 2, transferred to Belitsky minimal medium and treated with 100 μM NaOCl for 60 and 90 min. Cytoplasmic proteins were prepared and subjected to non-reducing BSH-specific Western blot analysis using the polyclonal rabbit anti-BSH antiserum as described previously (Chi et al., 2013). The de-bacillithiolation reactions with purified GapDH-SSB and the BrxA/BSH/YpdA/NADPH pathway were also subjected to non-reducing BSH-specific Western blots.

### Brx-roGFP2 and Tpx-roGFP2 biosensor measurements

*S. aureus* COL, *ypdA* and *brxAB* mutant strains expressing the Brx-roGFP2 and Tpx-roGFP2 biosensor plasmids were grown in LB and used for measurements of the biosensor oxidation degree (OxD) along the growth curves and after injection of the oxidants H_2_O_2_ and NaOCl as described previously (Loi et al., 2017). The fully reduced and oxidized control samples of Tpx-roGFP2 expression strains were treated with 15 mM DTT and 20 mM cumene hydroperoxide, respectively. The Brx-roGFP2 and Tpx-roGFP2 biosensor fluorescence emission was measured at 510 nm after excitation at 405 and 488 nm using the CLARIOstar microplate reader (BMG Labtech). The OxD of the Brx-roGFP2 and Tpx-roGFP2 biosensors was determined for each sample and normalized to fully reduced and oxidized controls as described (Loi et al., 2017).

### Biochemical assays for NADPH-dependent BSSB reduction by YpdA and de-bacillithiolation of GapDH-SSB using the BrxA/BSH/YpdA electron pathway *in vitro*

Before the activity assays, the purified BrxA, YpdA and YpdAC14A proteins were pre-reduced with 10 mM DTT followed by DTT removal with Micro Biospin 6 columns (Biorad). For the biochemical activity assays of the specific BSSB reductase activity, 12.5 μM of purified YpdA and YpdAC14A proteins were incubated with 40 μM BSSB or 40 μM coenzyme A disulfide and 500 μM NADPH in 20 mM Tris, 1.25 mM EDTA, pH 8.0. NADPH consumption of YpdA and YpdAC14A was measured immediately after the start of the reaction as absorbance change at 340 nm using the Clariostar microplate reader. The NADPH-dependent BrxA/BSH/YpdA electron pathway was reconstituted *in vitro* for de-bacillithiolation of GapDH-SSB. About 60 μM of purified GapDH was *S*-bacillithiolated with 600 μM BSH in the presence of 6 mM H_2_O_2_ for 5 min. Excess of BSH and H_2_O_2_ were removed with Micro Biospin 6 columns, which were equilibrated with 20 mM Tris, 1.25 mM EDTA, pH 8.0. Before starting the de-bacillithiolation assay using the BrxA/BSH/YpdA electron pathway, 2.5 μM GapDH-SSB was incubated with 12.5 μM BrxA, 40 μM BSH and 500 μM NADPH in 20 mM Tris, 1.25 mM EDTA, pH 8.0 at room temperature for 30 min. Next, 12.5 μM YpdA or YpdAC14A proteins were added to the reaction mix at 30°C for 8 min and NADPH consumption was measured at 340 mm. The biochemical activity assays were performed in four replicate experiments.

### Infection assays with murine macrophage cell line J-774A.1

The murine cell line J774A.1 was cultivated in Iscove’s modified Dulbecco MEM medium (Biochrom) with 10% heat inactivated foetal bovine serum (FBS) and used for *S. aureus* infection assays as described (Loi et al., 2018b). Macrophages were infected with *S. aureus* cells at a multiplicity of infection (MOI) of 1:25. One hour after infection, the cell culture medium was replaced and 150 μg/ml gentamycin was added for one hour to kill extracellular bacteria and to stop the uptake of *S. aureus*. The *S. aureus* cells were harvested at 2, 4 and 24 hours post infection. To determine the percentage of surviving *S. aureus* cells, infected macrophages were lysed with 0.1 % Triton X-100 and the supernatant of internalized bacteria was plated on brain heart infusion (BHI) agar plates. The CFUs were analyzed after incubation for 24-36 h at 37°C (Loi et al., 2018b).

### Statistical analyses

Statistical analysis of growth and survival assays was performed using the Student's unpaired two-tailed *t*-test by the graph prism software. The statistics of the J-774.1 macrophage infection assays was calculated using the one-way ANOVA and Tukey’s multiple comparisons post hoc test by the graph prism software. The results of the statistical tests are included in the figure legends.

## RESULTS

### Transcription of *ypdA*, *brxA* and *brxB* is induced under disulfide stress by diamide and NaOCl in *S. aureus* COL

The bacilliredoxins BrxA (SACOL1464) and BrxB (SACOL1558) of *S. aureus* share an unusual CGC active site and are highly conserved across BSH-producing firmicutes **(Fig. S1)** (Gaballa et al., 2014). The pyridine nucleotide disulfide oxidoreductase YpdA (SACOL1520) belongs to the FAD/NAD(P)-binding domain superfamily (IPR036188) and was annotated as putative BSSB reductase due to its phylogenetic co-occurence with the BSH biosynthesis enzymes in firmicutes **(Fig. S1)** (Gaballa et al., 2010). We used Northern blot analysis to investigate whether transcription of *brxA*, *brxB* and *ypdA* is co-regulated and up-regulated under thiol-specific stress conditions, such as 0.5 mM methylglyoxal, 0.75 mM formaldehyde, 1 mM NaOCl, 10 mM H_2_O_2_, 2 mM diamide and 45 μM methylhydroquinone (Fig. 1C). The *brxA* gene is co-transcribed with *SACOL1465-66-67* in a 2 kb operon and *brxB* is located in the 1.6 kb *SACOL1557-brxB-SACOL1559* operon. The Northern blot results revealed significant basal transcription of the *brxA, brxB* and *ypdA* genes and operons in the control, and strong induction under disulfide stress provoked by NaOCl and diamide. Of note, the *brxB* operon was stronger induced under disulfide stress compared to the *brxA* operon (Fig. 1C). No up-regulation of the *brxA, brxB* and *ypdA* specific mRNAs was detected upon H_2_O_2_, aldehyde and quinone stress. The co-regulation of BrxA/B and YpdA under disulfide stress suggests that they act in the same pathway to regenerate *S*-bacillithiolated proteins under NaOCl stress upon recovery from oxidative stress.

### The BSSB level is significantly increased and the BSH/BSSB ratio is decreased in the *S. aureus ypdA* mutant

To investigate the physiological role of BrxA/B and YpdA under oxidative stress and in BSH redox homeostasis, we constructed *brxAB* and *ypdA* deletion mutants. Using HPLC thiol metabolomics, the intracellular levels of BSH and BSSB were determined in the *brxAB* and *ypdA* mutants under control and NaOCl stress after monobromobimane derivatisation of LMW thiols and disulfides. In the *S. aureus* COL wild type, a BSH level of 1.6-1.9 μmol/g raw dry weight (rdw) was determined, which was not significantly different in the *ypdA* and *brxAB* mutants (Fig. 2A). Exposure of *S. aureus* to 2 mM NaOCl stress caused a 5-6-fold decreased intracellular BSH level in the wild type, *ypdA* and *brxAB* mutants (Fig. 2A). The level of BSSB was similar in control and NaOCl-treated cells of the wild type and *brxAB* mutant (∼0.05 μmol/g rdw) (Fig. 2B). Most interestingly, the *ypdA* mutant showed a significantly 2-fold increased BSSB level under control and NaOCl stress compared to the wild type (Fig. 2B). Thus, the BSH/BSSB ratio is ∼2-3-fold decreased in the *ypdA* mutant under control and NaOCl relative to the parent (Fig. 2C). The increased BSSB levels and the decreased BSH/BSSB redox ratio in the *ypdA* mutant could be restored to wild type levels in the *ypdA* complemented strain. In addition, a significantly 1.5-fold increased cysteine level was measured in the *ypdA* mutant under NaOCl stress, but no changes in the level of cystine **(Fig. S2AB)**. The cysteine levels could be also restored to wild type level in the *ypdA* complemented strain. These results indicate that YpdA is important to maintain the reduced level of BSH under control and NaOCl stress, while the bacilliredoxins BrxA/B are dispensible for the cellular BSH/BSSB redox balance during the growth and under oxidative stress in *S. aureus*.

**Fig. 2.**
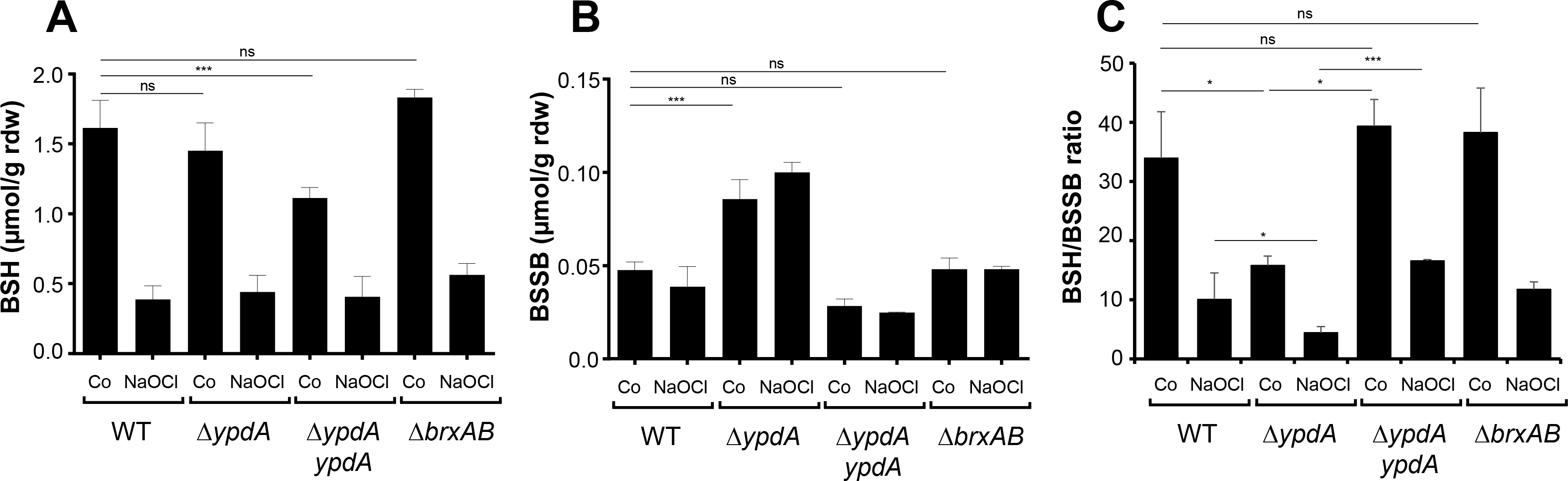
The BSSB level is strongly increased and the BSH/BSSB redox ratio is decreased in the *S. aureus* COL *ypdA* mutant under control and NaOCl stress. The levels of BSH **(A)** and BSSB **(B)** were determined in *S. aureus* COL wild type (WT), the *ypdA* and *brxAB* deletion mutants as well as in the *ypdA* complemented strain before (Co) and 30 min after treatment with 2 mM NaOCl stress. LMW thiols and disulfides were labelled with monobromobimane and measured using HPLC-thiol-metabolomics. **(C)** The BSH/BSSB ratio is significantly 2-fold decreased in the *ypdA* mutant compared to the WT and the *ypdA* complemented strain. The BSH and BSSB levels of the *brxAB* mutant are comparable to the wild type. Mean values are presented and error bars indicate the standard deviation of four biological replicates. ^ns^p > 0.05; **p ≤ 0.01 and ***p ≤ 0.001.

### The *S. aureus ypdA* mutant is impaired to regenerate the reduced BSH redox potential and to detoxify H_2_O_2_ under oxidative stress

Next, we applied the Brx-roGFP2 biosensor to monitor the changes in the BSH redox potential (*E*_BSH_) in *S. aureus* COL wild type, the *ypdA* and *brxAB* mutants (Loi et al., 2017). Measurements of the Brx-roGFP2 oxidation degree (OxD) along the growth in LB medium did not reveal notable differences in the basal level of *E*_BSH_ between wild type, *ypdA* and *brxAB* mutant strains **(Fig. S3AB)**. Thus, we monitored changes in *E*_BSH_ in *ypdA* and *brxAB* mutants after exposure to sub-lethal doses of 100 μM NaOCl and 100 mM H_2_O_2_ to identify functions for BrxAB or YpdA under oxidative stress. The Brx-roGFP2 biosensor was strongly oxidized under NaOCl and H_2_O_2_ stress in the wild type, *ypdA* and *brxAB* mutants (Fig. 3A-D). Regeneration of reduced *E*_BSH_ occurred within two hours in the wild type and *brxAB* mutants. However, the *ypdA* mutant was significantly impaired to recover the reduced state of BSH and the OxD remained high even after two hours of NaOCl and H_2_O_2_ stress (Fig. 3AC). Of note, the defect of the *ypdA* mutant to restore reduced *E*_BSH_ was reproducible with both oxidants, H_2_O_2_ and NaOCl, supporting the important role of YpdA to reduce BSSB during the recovery phase of cells from ROS stress.

**Fig. 3.**
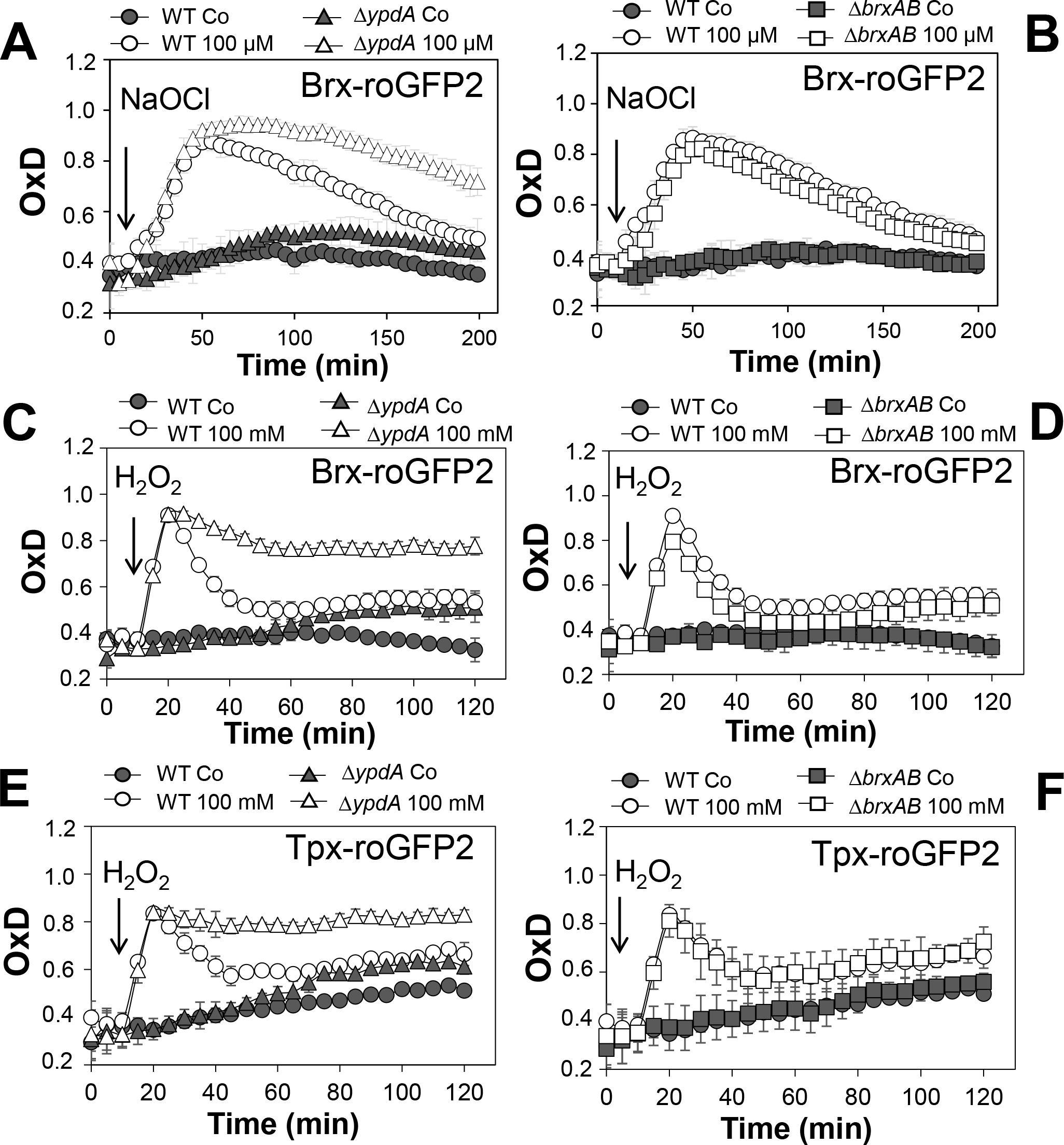
Brx-roGFP2 and Tpx-roGFP2 biosensor measurements indicate that the *S. aureus ypdA* mutant is impaired to regenerate *E* BSH and to detoxify H_2_O_2_ during recovery from oxidative stress. **(A-D)** Response of Brx-roGFP2 to 100 μM NaOCl and 100 mM H_2_O_2_ stress in *S. aureus* COL wild type (WT), the *ypdA* **(A, C)** and *brxAB* **(B, D)** mutants. The BSH redox potential is restored in the WT and in the *brxAB* mutant upon recovery from oxidative stress, **(B, D)**, but the *ypdA* mutant is unable to restore the reduced BSH redox potential in the recovery phase. **(E, F)** Response of the Tpx-roGFP2 biosensor under 100 mM H_2_O_2_ stress in the *S. aureus* COL WT, the *ypdA* and *brxAB* mutants. The Tpx-roGFP2 biosensor responds strongly to 100 mM H_2_O_2_ and is reduced after 60 min of H_2_O_2_ exposure in the WT and *brxAB* mutants. However, the *ypdA* mutant is impaired in detoxification of H_2_O_2_ during the recovery phase. In all graphs, mean values are presented and error bars indicate the standard deviation of three biological replicates.

We further hypothesized that the *ypdA* mutant is defective for H_2_O_2_ detoxification due to its increased BSSB levels. To analyse the kinetics of H_2_O_2_ detoxification in the *ypdA* mutant, we constructed a genetically encoded H_2_O_2_-specific Tpx-roGFP2 biosensor. First, we verified that Tpx-roGFP2 showed the same ratiometric changes of the excitation spectrum in the fully reduced and oxidized state *in vitro* and *in vivo* as previously measured for Brx-roGFP2 **(Fig. S4AB)**. Tpx-roGFP2 was shown to respond strongly to low levels of 0.5-1 μM H_2_O_2_ *in vitro* and was fully oxidized with 100 mM H_2_O_2_ inside *S. aureus* COL wild type cells indicating the utility of the probe to measure H_2_O_2_ detoxification kinetics in *S. aureus* **(Fig. S4CD)**. Measurements of Tpx-roGFP2 oxidation along the growth in LB medium revealed a similar high OxD of ∼0.5-0.6 in the wild type, *brxAB* and *ypdA* mutant strains **(Fig. S3CD)**. The absence of BrxA/B or YpdA did not affect the biosensor OxD under non-stress conditions, which further provides evidence for roles under oxidative stress. Thus, we monitored the H_2_O_2_ response of Tpx-roGFP2 and the kinetics of H_2_O_2_ detoxification in the *ypdA* and *brxAB* mutants. Interestingly, Tpx-roGFP2 showed a similar response to 100 mM H_2_O_2_ in all strains, but the *ypdA* mutant was significantly impaired in H_2_O_2_ detoxification compared to the wild type (Fig. 3EF). These results clearly confirmed that the *ypdA* mutant is defective to recover from oxidative stress due to its higher BSSB level resulting in an oxidized BSH redox potential as revealed using Brx-roGFP2 and thiol-metabolomics studies.

### *S*-bacillithiolation of GapDH is not affected in the *ypdA* and *brxAB* mutants

In *S. aureus*, the glyceraldehyde dehydrogenase GapDH was previously identified as most abundant *S*-bacillithiolated protein under NaOCl stress that is visible as major band in BSH-specific non-reducing Western blots (Imber et al., 2018a). Since GapDH activity could be recovered with purified BrxA *in vitro* previously (Imber et al., 2018a), we analyzed the pattern of GapDH *S*-bacillithiolation in the *brxAB* and *ypdA* mutants *in vivo*. However, the amount of *S*-bacillithiolated GapDH was similar after 100 μM NaOCl stress between wild type, *brxAB* and *ypdA* mutant cells (Fig. 4). This indicates that the absence of the BrxAB/YpdA pathway does not affect the level of *S*-bacillithiolation of GapDH under NaOCl stress.

**Fig. 4.**
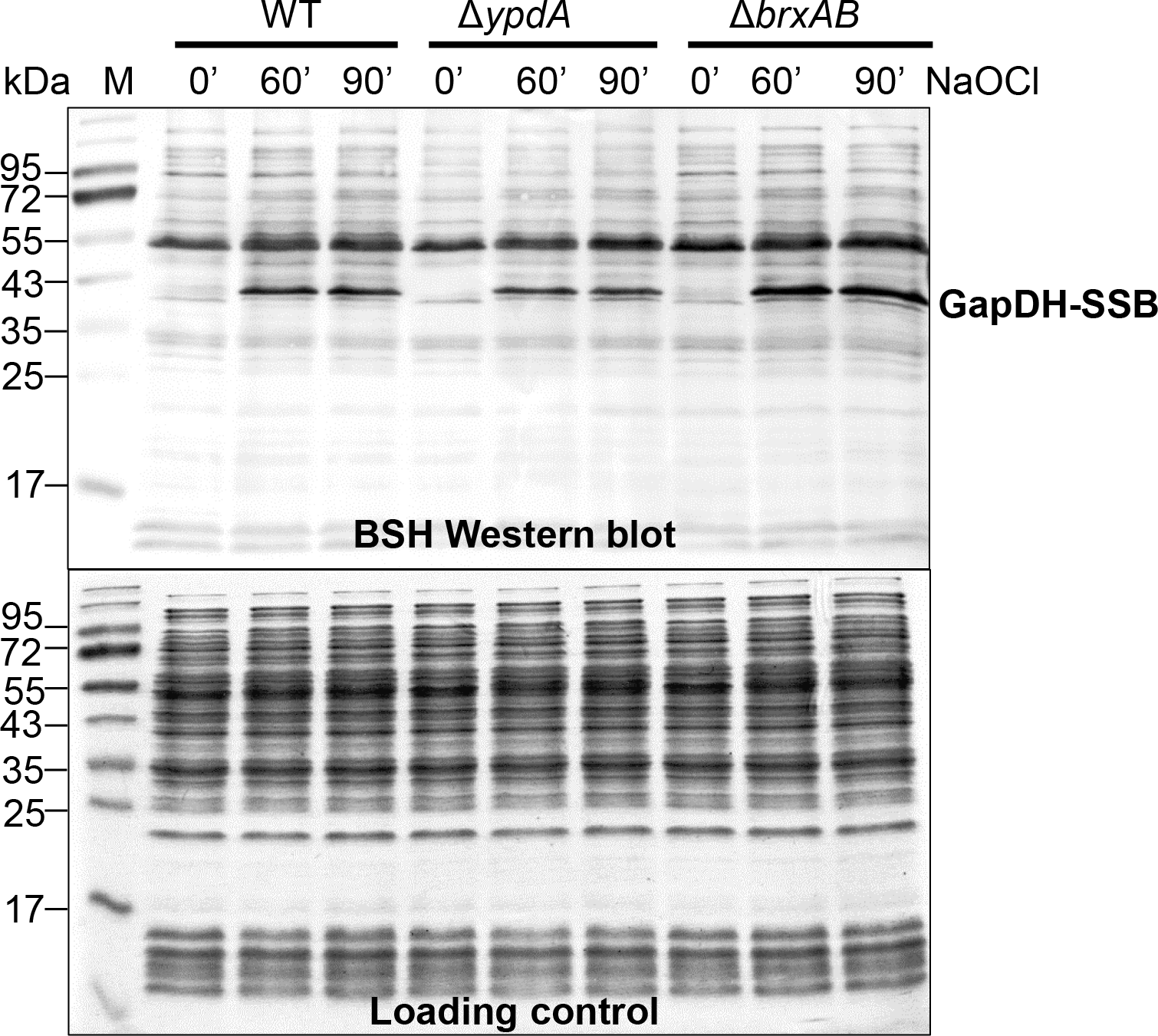
Protein *S*-bacillithiolation of GapDH is not affected in the *ypdA* or *brxAB* mutants as revealed by non-reducing BSH Western blots. The prominent GapDH-SSB band is visible in the cell extracts of NaOCl-treated *S. aureus* cells using non-reducing BSH Western blots. The amount of GapDH-SSB is similar in the WT, *ypdA* and *brxAB* mutant cells. The SDS PAGE loading control is shown at the bottom for comparison. Representative Western blots and loading controls of 3 biological replicates are shown.

### The bacilliredoxins BrxA/B and the putative BSSB reductase YpdA are important for growth and survival under oxidative stress and macrophage infections

Next, we analyzed the physiological role of the BrxA/B/YpdA pathway for growth and survival of *S. aureus* under H_2_O_2_ and NaOCl stress. The growth of the *ypdA* and *brxAB* mutants in RPMI medium without stress exposure was comparable to the wild type (Fig. 5AC). Interestingly, both *brxAB* and *ypdA* mutants were significantly delayed in growth after exposure to sub-lethal amounts of 1.5 mM NaOCl, but no growth delay was observed with sub-lethal 10 mM H_2_O_2_ (Fig. 5AC, 6AB). Determination of viable counts revealed significantly ∼2-fold decreased survival rates of both *brxAB* and *ypdA* mutants after exposure to lethal doses of 3.5 mM NaOCl and 40 mM H_2_O_2_ relative to the wild type (Fig. 5FG, 6CD). These oxidant sensitive growth and survival phenotypes of the *brxAB* and *ypdA* mutants could be restored back to wild type levels by complementation with *brxA* and *ypdA* respectively (Fig. 5BDFG, 6CD). However, complementation of the *brxAB* mutant with *brxB* did not restore the growth and viability of the wild type under NaOCl stress (Fig. 5EG). Moreover, the *brxAB ypdA* triple mutant displayed the same sensitivity as the *brxAB* mutant to 40 mM H_2_O_2_ and 3 mM NaOCl indicating that BrxA and YpdA function in the same pathway for reduction of *S*-bacillithiolated proteins (Fig. 6D**, S6C)**.

**Fig. 5.**
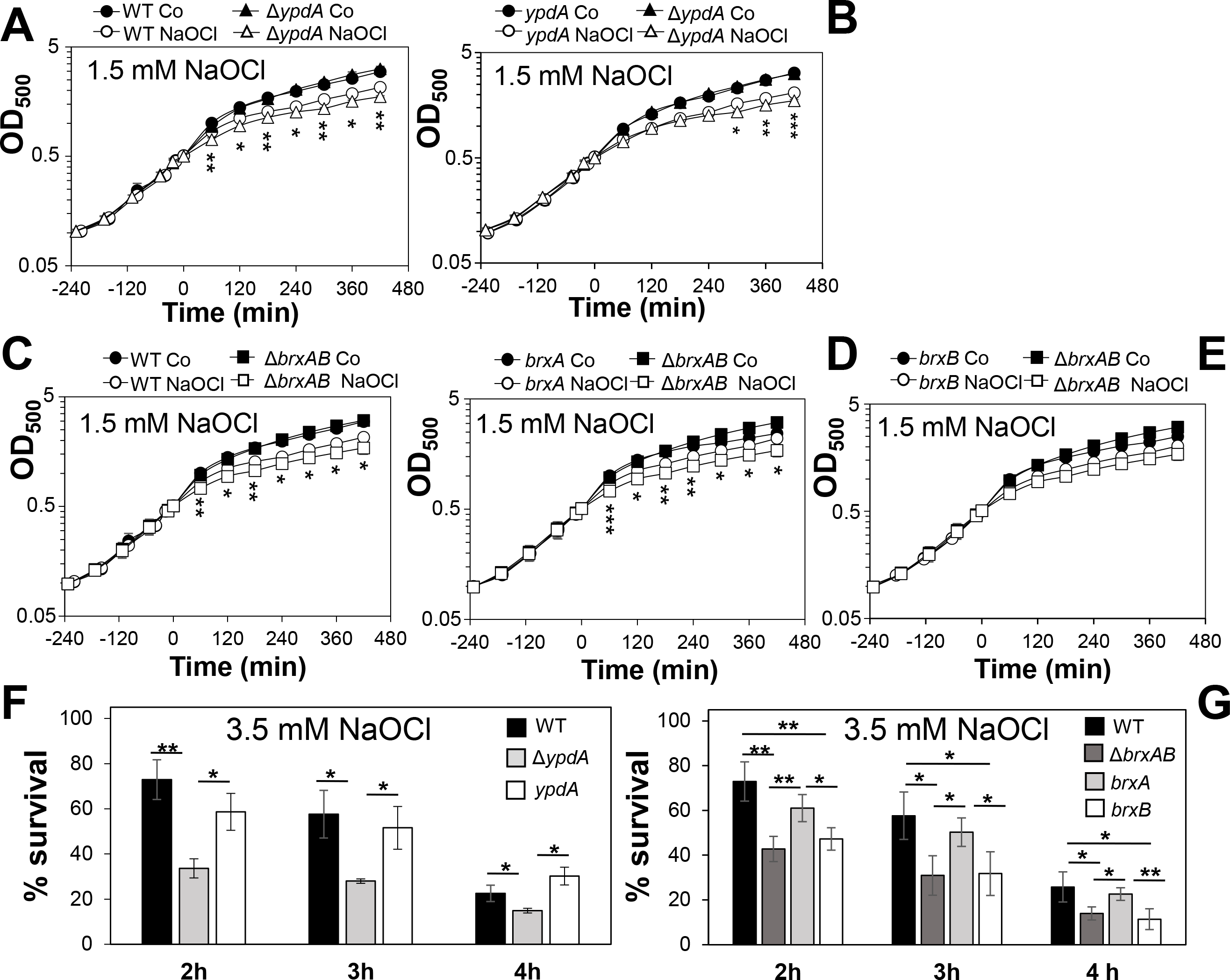
The *S. aureus ypdA* and *brxAB* mutants are more sensitive under NaOCl stress. **(A-E)** Growth curves of *S. aureus* COL WT, the *ypdA* and *brxAB* mutants as well as the *brxA* and *brxB* complemented strains after exposure to a sub-lethal 1.5 mM NaOCl stress. The cells were grown in RPMI medium until an OD_500_ of 0.5 and treated with NaOCl. **(F,G)** Survival rates were determined as CFUs for *S. aureus* COL WT, *ypdA* and *brxAB* mutants and *brxA, brxB* complemented strains at 2, 3 and 4 hours after treatment with 3.5 mM NaOCl. Survival of the untreated control was set to 100%. Mean values are presented of 3-4 biological replicates. Error bars indicate the standard deviation and the statistics was calculated using a Student's unpaired two-tailed t-test by the graph prism software. Symbols are: ^ns^ p > 0.05, *p ≤ 0.05 and **p ≤ 0.01.

**Fig. 6.**
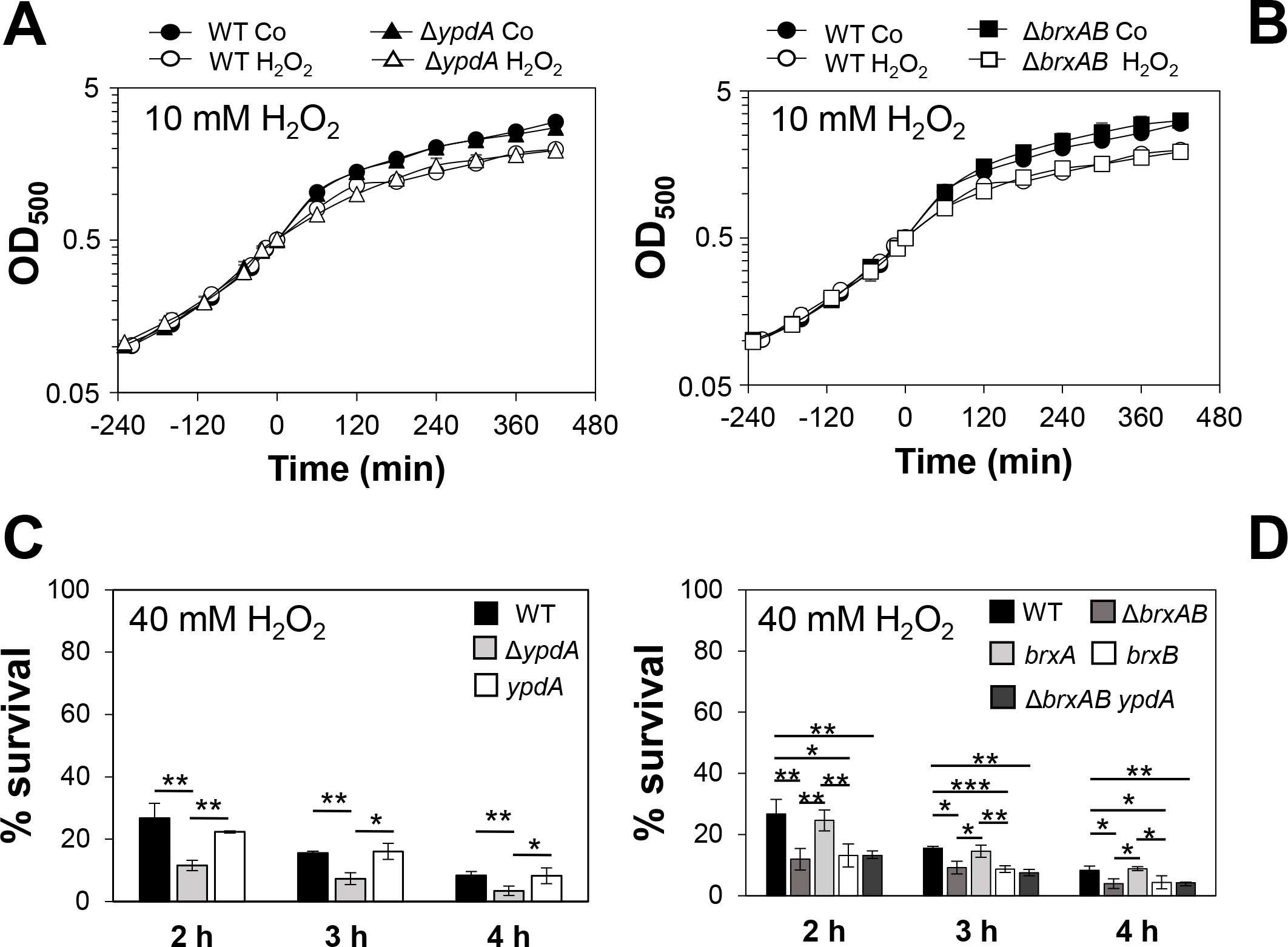
The *S. aureus ypdA* and *brxAB* mutants show increased sensitivities under H_2_O_2_ stress. **(A, B)** Growth curves of *S. aureus* COL WT, the *ypdA* and *brxAB* mutants and *brxA, brxB* complemented strains after exposure to a sub-lethal 10 mM H_2_O_2_ stress. The cells were grown in RPMI medium until an OD_500_ of 0.5 and treated with H_2_O_2_. **(C, D)** Survival rates were determined as CFUs for *S. aureus* COL WT, *ypdA*, *brxAB* and *brxAB ypdA* mutants and *brxA, brxB* complemented strains at 2, 3 and 4 hours after treatment with 40 mM H_2_O_2_. Survival of the untreated control was set to 100%. Mean values are presented from 3-5 biological replicates. Error bars indicate the standard deviation and the statistics was calculated using a Student’s unpaired two-tailed t-test by the graph prism software. Symbols are : ^ns^p > 0.05, *p≤ 0.05 and **p ≤ 0.01.

To investigate the function of the BrxA/B/YpdA pathway under infection-relevant conditions, we measured the intracellular survival of the *brxAB* and *ypdA* mutants in phagocytosis assays inside murine macrophages of the cell line J-774A.1, as previously (Loi et al., 2018b). The viable counts (CFUs) of internalized *S. aureus* cells were determined at 2, 4 and 24 hours post infection (p.i.) of the macrophages. The number of surviving cells decreased to 21.3% at 24 hours p.i. for the *S. aureus* COL wild type, but more strongly to 11.4% and 10.2% for the *ypdA* and *brxAB* mutants (Fig. 7AC). Thus, the number of viable counts was significantly ∼2-fold lower for both *brxAB* and *ypdA* mutants at 24 h p.i. compared to the wild type. These sensitive phenotypes of the *ypdA* and *brxAB* mutants under macrophage infections could be restored to 80% of wild type levels after complementation with plasmid-encoded *ypdA* or *brxA*, respectively (Fig. 7BD). However, complementation with *brxB* did not restore the survival defect of the *brxAB* mutant, pointing again to the major role of BrxA in this pathway.

**Fig. 7.**
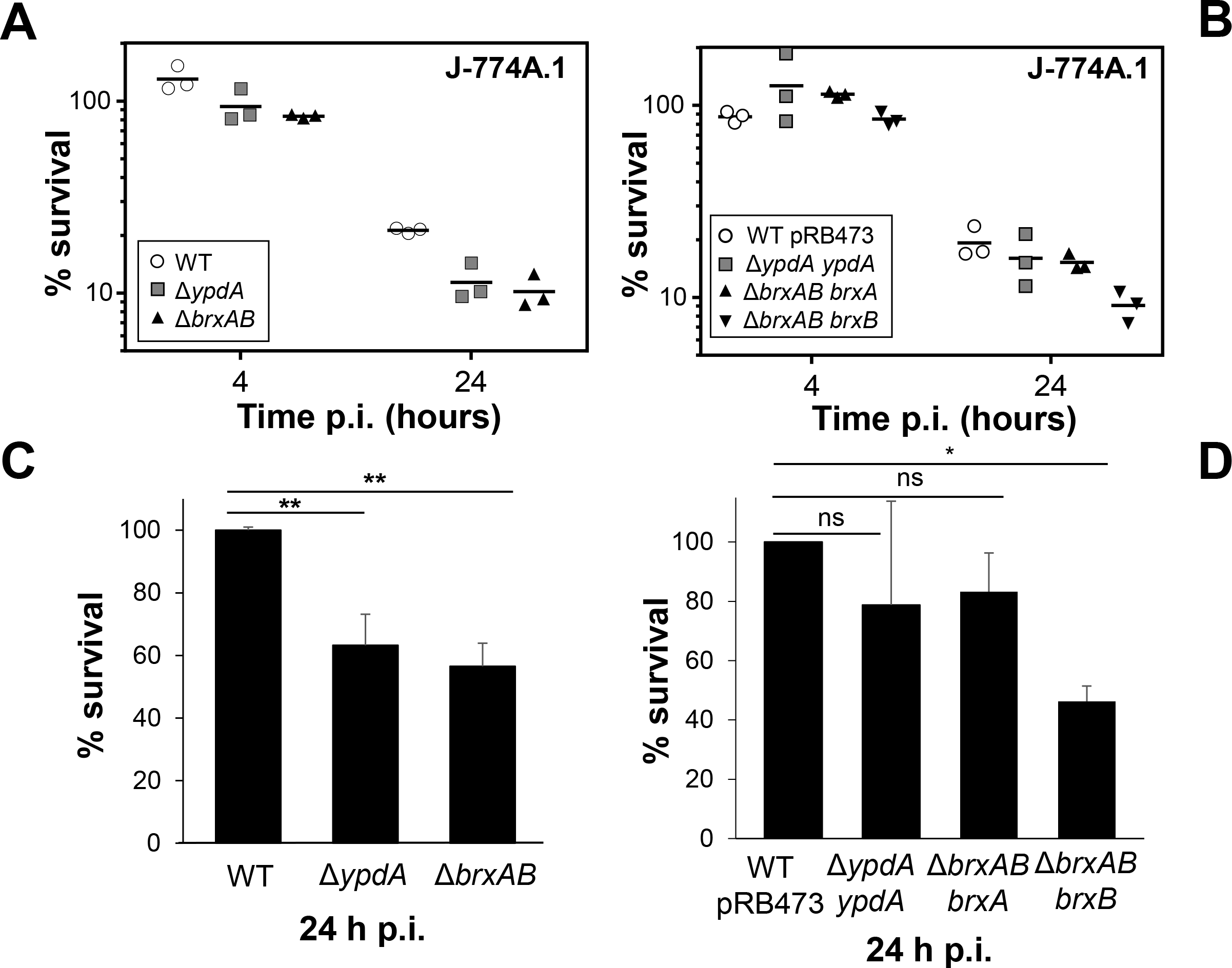
YpdA and BrxA/B promote the intracellular survival of *S. aureus* inside murine macrophages during infections. **(A, B)** The survival of *S. aureus* WT*, ypdA* and *brxAB* mutants and complemented strains was analyzed 2, 4 and 24 h post infection of the murine macrophage cell line J-774A.1 by CFU counting. The percentages in survival of the *ypdA* and *brxAB* mutants and complemented strains were calculated after 4 and 24 hours in relation to the 2 hour time point, which was set to 100%. **(C, D)**. The average percentage in survival was calculated for *ypdA* and *brxAB* mutants **(C)** and complemented strain **(D)** in relation to the WT and WT with empty plasmid pRB473, which were set to 100%. Error bars represent the SEM and the statistics were calculated using one-way ANOVA and Tukey’s multiple comparisons post hoc test using the graph prism software (*p* = 0.0050 for WT/∆*ypdA*, *p* = 0.0022 for WT/∆*brxAB* and *p* = 0.026 for WT pRB473/∆*brxAB brxB*). Symbols: ^ns^p > 0.05; *p ≤ 0.05 and **p ≤ 0.01.

Taken together, our results revealed that the bacilliredoxin BrxA and the putative BSSB reductase YpdA are required for improved survival of *S. aureus* inside macrophages to resist the oxidative burst. Our data suggest that BrxA and YpdA act together in the BrxA/BSH/YpdA pathway to regenerate *S*-bacillithiolated proteins and to restore the BSH redox potential upon recovery from oxidative stress during infections.

### The flavin disulfide reductase YpdA functions in BSSB reduction and de-bacillithiolation of GapDH-SSB in the BrxA/BSH/YpdA electron transfer assay *in vitro*

Next, we aimed to analyze the catalytic activity of purified YpdA in a NADPH-coupled assay with BSSB as substrate *in vitro*. The His-tagged YpdA protein was purified as yellow coloured enzyme and the UV-visible spectrum revealed the presence of the FAD co-factor indicated by the two absorbance peaks at 375 and 450 nm **(Fig. S5)**. Incubation of YpdA protein with BSSB resulted in significant and fast consumption of NADPH as measured by a rapid absorbance decrease at 340 nm (Fig. 8A). Only little NADPH consumption was measured with YpdA alone in the absence of the BSSB substrate supporting previous finding that YpdA consumes NADPH alone (Mikheyeva et al., 2019). However, in our assays, BSSB significantly enhanced NADPH consumption by YpdA compared to the control reaction without BSSB. No increased NADPH consumption was measured with coenzyme A disulphide (CoAS_2_) as substrate indicating the specificity of YpdA for BSSB (Fig. 8A). In addition, we investigated the role of the conserved Cys14 of YpdA for the BSSB reductase activity in the NADPH-coupled assay. NADPH-consumption of YpdAC14A upon BSSB reduction was much slower and similar to the control reaction of YpdA and YpdAC14A without BSSB (Fig. 8B).

**Fig. 8.**
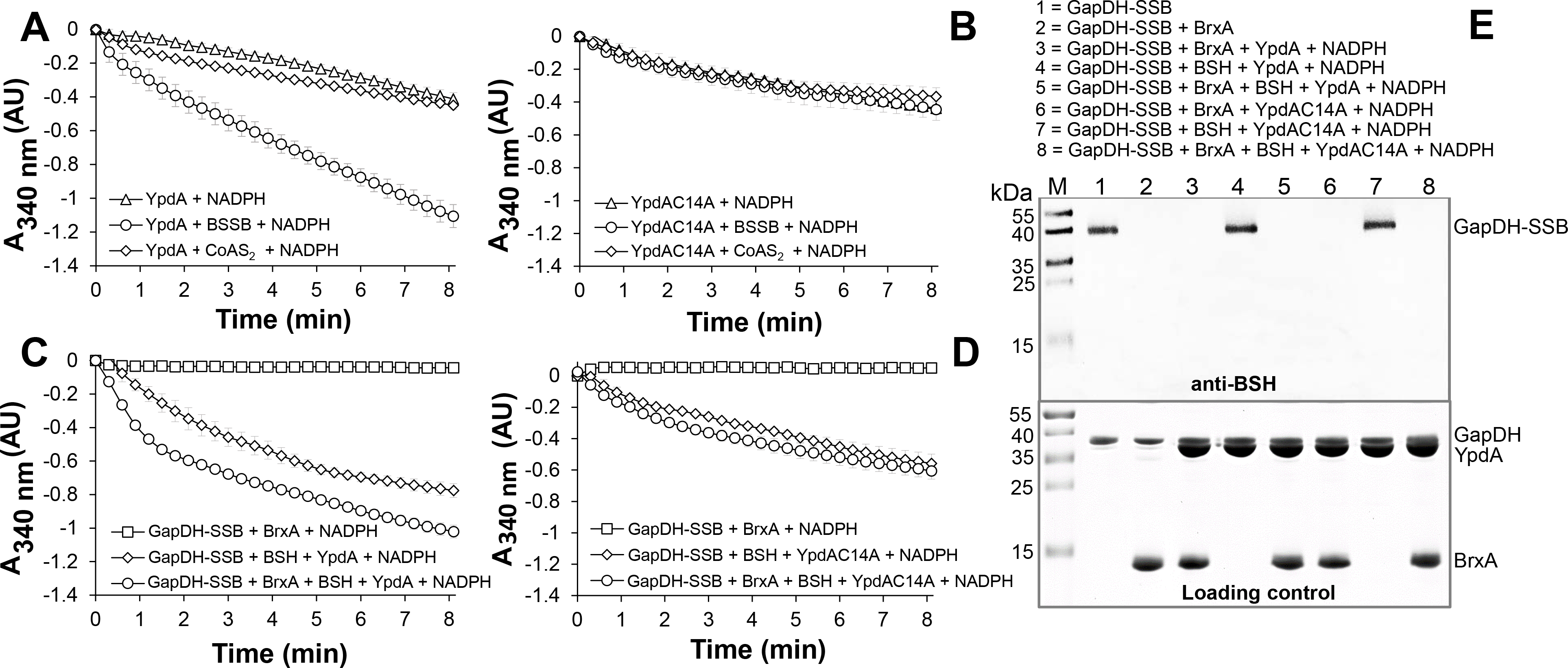
YpdA functions as BSSB reductase in the NADPH-coupled BrxA/BSH/YpdA electron pathway for de-bacillithiolation of GapDH-SSB *in vitro*. **(A)** Purified YpdA is able to reduce BSSB back to BSH with electrons from NADPH as measured by the absorbance change at 340 nm. Only little NADPH consumption was measured with YpdA alone in the absence of BSSB, with coenzyme A disulfide (CoAS_2_) **(A)** or in the YpdAC14A mutant **(B)** indicating the function of the conserved Cys14 as active site of YpdA. **(C)** NADPH consumption of YpdA was measured in the coupled BrxA/BSH/YpdA de-bacillithiolation assay for reduction of GapDH-SSB. While fast NADPH consumption was measured upon de-bacillithiolation of GapDH-SSB with BrxA/BSH/YpdA **(C)**, the reaction was much slower with BrxA/BSH/YpdAC14A **(D)**. The coupled assays were conducted with 2.5 μM GapDH-SSB, 12.5 μM BrxA, 40 μM BSH and 500 μM NADPH in 20 mM Tris, 1.25 mM EDTA, pH 8.0. After 30-min incubation at room temperature, 12.5 μM YpdA or YpdAC14A proteins were added to the reaction mix and NADPH consumption was monitored at 340 nm as a function of time. Mean values and SEM of four independent experiments are shown. **(E)** The de-bacillithiolation of GapDH-SSB is catalyzed by BrxA or the complete BrxA/BSH/YpdA pathway, but not by YpdA alone as shown by non-reducing BSH-specific Western blots. The loading controls are shown below as Coomassie-stained SDS-PAGE with the same de-bacillithiolation reactions of GapDH-SSB as in the BSH-blot.

Our *in vivo* data support that YpdA and BrxA act together in the BrxA/BSH/YpdA de-bacillithiolation pathway. Thus, we analysed NADPH-consumption by the BrxA/BSH/YpdA electron pathway in de-bacillithiolation of GapDH-SSB *in vitro*. The de-bacillithiolation assays revealed fast NADPH consumption in the complete BrxA/BSH/YpdA coupled assays (Fig. 8C). NADPH consumption by YpdA was slower in the absence of BrxA and might be caused by residual BSSB in the BSH samples. The control reaction of GapDH-SSB with BrxA did not consumed NADPH and only little NADPH consumption was measured with BrxA, BSH and the YpdAC14A mutant protein in de-bacillithiolation of GapDH-SSB (Fig. 8D).

In addition, BSH-specific non-reducing Western blots were used to investigate if BrxA and the complete BrxA/BSH/YpdA pathway catalyse de-bacillithiolation of GapDH-SSB (Fig. 8E). The BSH-blots showed that BrxA is sufficient for de-bacillithiolation of GapDH-SSB, since all reactions of GapDH-SSB with BrxA lead to complete de-bacillithiolation with and without YpdA or YpdAC14A plus NADPH. However, the reactions of GapDH-SSB with YpdA/NADPH alone did not lead to reduction of GapDH-SSB, indicating the main role of BrxA in de-bacillithiolation while YpdA functions in regeneration of BSH in the BrxA/BSH/YpdA/NADPH redox cycle.

In conclusion, our biochemical assays revealed that YpdA functions as BSSB reductase in an NADPH coupled assay. Cys14 of YpdA is important for the BSSB reductase activity *in vitro*. Thus, YpdA facilitates together with BrxA the reduction of *S*-bacillithiolated GapDH in the BrxA/BSH/YpdA redox pathway upon recovery from oxidative stress.

## Discussion

In this work, we have studied the role of the bacilliredoxins BrxA/B and the BSSB reductase YpdA in the defense of *S. aureus* against oxidative stress. Transcription of *brxA*, *brxB* and *ypdA* is strongly upregulated under disulfide stress, provoked by diamide and NaOCl. About 2-4 fold increased transcription of *ypdA, brxA* and *brxB* was previously found under H_2_O_2_, diamide, NaOCl, AGXX^®^ and after exposure to azurophilic granule proteins in *S. aureus* (Palazzolo-Ballance et al., 2008;Posada et al., 2014;Mäder et al., 2016;Loi et al., 2018a;Loi et al., 2018b). The elevated transcription of *brxA, brxB* and *ypdA* under disulfide stress correlated with the up-regulation of the *bshA*, *bshB* and *bshC* genes for bacillithiol biosynthesis in *S. aureus* and *B. subtilis* (Chi et al., 2011;Nicolas et al., 2012;Loi et al., 2018a;Loi et al., 2018b). The *bshA, bshB* and *bshC* genes and operons are under control of the disulfide stress-specific Spx regulator in *B. subtilis*, which controls a large regulon for thiol-redox homeostasis (Gaballa et al., 2013). Thus, genes for BSH biosynthesis and the BrxA/B/YpdA pathway might be also regulated by Spx in *S. aureus*.

The co-regulation of BrxA/B and YpdA under disulfide stress points to their function in the same pathway in *S. aureus*. HOCl, diamide and AGXX^®^ were shown to cause a strong disulfide stress response in the transcriptome and protein *S*-bacillithiolation in the proteome of *S. aureus* (Imber et al., 2018a;Loi et al., 2018a;Loi et al., 2018b). Thus, the BrxA/B and YpdA redox enzymes are up-regulated under conditions of protein *S*-bacillithiolations, connecting their functions to the de-bacillithiolation pathway. We could show here that NaOCl stress leads to 3-4-fold depletion of the cellular pool of reduced BSH in the *S. aureus* COL wild type, which was not accompanied by an enhanced BSSB level. Most probably, the increased BSSB level under NaOCl stress was used for protein *S*-bacillithiolation (Imber et al., 2018a).

The BSH/BSSB redox ratio of *S. aureus* wild type cells was determined as ∼35:1 under control conditions and decreased 3-fold to 10:1 under NaOCl. Similarly, the GSH/GSSG redox ratio was determined in the range between 30:1 and 100:1 in the cytoplasm of *E. coli*, which is kept reduced by the NADPH-dependent glutathione reductase (Gor) (Hwang et al., 1995;Van Laer et al., 2013). In the *S. aureus brxAB* mutant, we also measured a 3-fold decrease of the BSH/BSSB ratio from control conditions (38:1) to NaOCl (12:1). However, the *ypdA* mutant showed a 2-fold enhanced BSSB level in control and NaOCl-treated cells, leading to a significantly decreased BSH/BSSB ratio under control (17:1) and NaOCl stress (5:1). Thus, our data indicate that BrxAB are dispensable for the BSH redox homeostasis, while YpdA is essential for BSSB reduction to maintain the reduced pool of BSH and a high BSH/BSSB ratio in *S. aureus*.

Brx-roGFP2 biosensor measurements provide further support that YpdA is the candidate BSSB reductase. The *ypdA* mutant was significantly impaired to restore reduced *E*_BSH_ during recovery from NaOCl and H_2_O_2_ stress. Moreover, application of the Tpx-roGFP2 biosensor revealed a delay in H_2_O_2_ detoxification in *ypdA* mutant cells during the recovery phase. These results clearly support the important role of YpdA as BSSB reductase particularly under oxidative stress to recover reduced *E*_BSH_ required for detoxification of ROS.

These *in vivo* data were further corroborated by biochemical activity assays of YpdA for BSSB reduction in a NADPH-coupled assay. While little NADPH consumption was measured in the presence of YpdA alone, BSSB significantly enhanced NADPH consumption, supporting the crucial role of YpdA as BSSB reductase *in vitro*. Further electron transfer assays revealed that YpdA functions together with BrxA and BSH in reduction of GapDH-SSB *in vitro*. Previous de-bacillithiolation assays have revealed regeneration of GapDH activity by BrxA *in vitro* (Imber et al., 2018a). Here, we confirmed that BrxA activity is sufficient for complete de-bacillithiolation of GapDH-SSB *in vitro*, while YpdA alone had no effect on the GapDH-SSB reduction. Thus, BrxA catalyzes reduction of *S*-bacillithiolated proteins and YpdA is involved in BSH regeneration in the complete BrxA/BSH/YpdA redox cycle.

The BSSB reductase activity of YpdA was shown to be dependent on the conserved Cys14, which is located in the glycine-rich Rossmann-fold NAD(P)H binding domain (GGGPC_14_G) (Bragg et al., 1997;Mikheyeva et al., 2019). Cys14 might be *S*-bacillithiolated by BSSB and reduced by electron transfer from NADPH via the FAD co-factor. Cys14 was previously identified as oxidized under NaOCl stress in the *S. aureus* redox proteome using the OxICAT method, further supporting its role as active site Cys and its S-bacillithiolation during the BrxA/BSH/YpdA catalytic cycle (Imber et al., 2018a). The catalytic mechanism of BSSB reduction via Cys14 of YpdA is an interesting subject of future studies.

Detailed phenotype analyses further revealed protective functions of the BrxA/BSH/YpdA redox pathway for growth and survival of *S. aureus* under oxidative stress *in vitro* and under macrophage infections *in vivo*. The *ypdA* and *brxAB* mutants were significantly impaired in growth and survival after exposure to sub-lethal and lethal doses of NaOCl and displayed survival defects under lethal H_2_O_2_. Moreover, the H_2_O_2_ and NaOCl-sensitivity and the defect to recover reduced *E*_BSH_ in the *brxAB ypdA* triple mutant was comparable with that of the *ypdA* mutant (Fig. 6D**, S6)**. These results clearly indicate that BrxA/B and YpdA function in the same de-bacillithiolation pathway.

Based on previous bacilliredoxin activity assays *in vitro*, both BrxA and BrxB should use a monothiol mechanism to reduce *S*-bacillithiolated client proteins, such as OhrR, GapDH and MetE in *B. subtilis* and *S. aureus* (Gaballa et al., 2014;Imber et al., 2018a). Most di-thiol glutaredoxins of *E. coli* (Grx1, Grx2 and Grx3) use the monothiol mechanism for de-glutathionylation of proteins (Lillig et al., 2008;Allen and Mieyal, 2012;Loi et al., 2015). In the monothiol mechanism, the nucleophilic thiolate of the Brx CGC motif attacks the *S*-bacillithiolated protein, resulting in reduction of the protein substrate and Brx-SSB formation. Brx-SSB is then recycled by BSH, leading to increased BSSB formation. YpdA reduces BSSB back to BSH with electrons from NADPH (Fig. 1B). The oxidation-sensitive phenotypes of *ypdA* and *brxAB* mutants could be complemented by plasmid-encoded *ypdA* and *brxA*, but not *brxB*, respectively. These results provide evidence for the function of the BrxA/BSH/YpdA de-bacillithiolation pathway using the monothiol-Brx mechanism in *S. aureus*.

Similar phenotypes were found for mutants lacking related redox enzymes of the GSH and mycothiol pathways in other bacteria. In *E. coli*, strains lacking the glutathione disulfide reductase and glutaredoxins are more sensitive under diamide and cumene hydroperoxide stress (Alonso-Moraga et al., 1987;Vlamis-Gardikas et al., 2002;Lillig et al., 2008). In *Mycobacterium smegmatis*, the mycoredoxin-1 mutant displayed an oxidative stress-sensitive phenotype (Van Laer et al., 2012). In *Corynebacterium glutamicum*, deficiency of the mycothiol disulfide reductase Mtr resulted in an oxidized mycothiol redox potential (Tung et al., 2019), and Mtr overexpression contributed to improved oxidative stress resistance (Si et al., 2016). Taken together, our results revealed that not only BSH, but also BrxA and YpdA are required for virulence and promote survival in infection assays inside murine macrophages.

In several human pathogens, such as *Streptococcus pneumoniae, Listeria monocytogenes, Salmonella* Typhimurium and *Pseudomonas aeruginosa*, LMW thiols or the glutathione disulfide reductase are required for virulence, colonization and to resist host-derived oxidative or nitrosative stress (Potter et al., 2012;Song et al., 2013;Reniere et al., 2015;Tung et al., 2018;Wongsaroj et al., 2018). *S. aureus* BSH deficient mutants showed decreased survival in murine macrophages and in human whole blood infections (Pöther et al., 2013;Posada et al., 2014). The virulence mechanisms might be related to a lack of BSH regeneration and decreased recovery of inactivated *S*-bacillithiolated proteins inside macrophages. Future studies should elucidate the targets for *S*-bacillithiolations that are reduced by the BrxA/BSH/YpdA pathway inside macrophages, increasing survival, metabolism or persistence under infections.

In summary, our results showed the importance of the BrxA/BSH/YpdA redox pathway to resist oxidative stress and macrophage infection in *S. aureus*. Through measurements of the BSH/BSSB redox ratio and *E*_BSH_, we provide evidence that the NADPH-dependent disulfide reductase YpdA regenerates BSH and restores reduced *E*_BSH_ upon recovery from oxidative stress in *S. aureus*. Finally, biochemical evidence for YpdA as BSSB reductase and for the role of BrxA/BSH/YpdA pathway in de-bacillithiolation was provided *in vitro*. The detailed biochemical mechanism of YpdA and the cross-talk of the Trx and Brx systems in de-bacillithiolation under oxidative stress and infections are subject of our future studies.

## ACKNOWLEDGEMENTS

We acknowledge support by the Open Access Publication Initiative of Freie Universität Berlin. This work was supported by an ERC Consolidator grant (GA 615585) MYCOTHIOLOME and grants from the Deutsche Forschungsgemeinschaft (AN746/4-1 and AN746/4-2) within the SPP1710, by the SFB973 project C08N and by the SFB/TR84 project B06 to H.A.

## LIST OF ABBREVIATIONS

BSH: bacillithiol
BSSB: bacillithiol disulfide
BrxA/B: bacilliredoxin A (YphP)/ bacilliredoxin B (YqiW)
CFUs: colony forming units
DTT: dithiothreitol
E_BSH_: bacillithiol redox potential
GapDH: glyceraldehyde 3-phosphate dehydrogenase
GSH: glutathione
GSSG: glutathione disulfide
Gor: glutathione disulfide reductase
Grx: glutaredoxin
HOCl: hypochlorous acid
LMW: low molecular weight
Mtr: mycothiol disulfide reductase
NaOCl: sodium hypochlorite
OD_500_: optical density at 500 nm
rdw: raw dry weight
RCS: reactive chlorine species
ROS: reactive oxygen species
YpdA: bacillithiol disulfide reductase

