## Supplemental Tables S1-S3 for "*Staphylococcus aureus* uses the bacilliredoxin (BrxAB)/ bacillithiol disulfide reductase (YpdA) redox pathway to defend against oxidative stress under infections"

**Table S1. Bacterial strains**

| Strain | Description | Reference |
|---|---|---|
| ***Escherichia coli*** | | |
| DH5α | F-φ80dlacZ Δ(lacZYA-argF) U169 deoRsupE44ΔlacU169 (f80lacZDM15) hsdR17 recA1 endA1 (rk- mk+) supE44gyrA96 thi-1 gyrA69 relA1 | [1] |
| BL21(DE3)*plys*S | F- ompT hsdS gal (rb- mb+) DE3(Sam7 Δnin5 lacUV5-T7 Gen1) | [1] |
| ***Staphylococcus aureus*** | | |
| RN4220 | restriction negative strain/MSSA cloning intermediate derived from 8325-4 | [2] |
| COL | Archaic HA-MRSA strain | [3] |
| COL Δ*ypdA* | COL *ypdA* deletion mutant | This study |
| COL Δ*brxA* | COL *brxA* deletion mutant | This study |
| COL Δ*brxAB* | COL *brxAB* double mutant | This study |
| COL Δ*brxAB ypdA* | COL *brxAB ypdA* triple mutant | This study |
| COL pRB473 | | [4] |
| COL pRB473-*brx-roGFP2* | | [4] |
| COL Δ*ypdA::*pRB473-*brx-roGFP2* | | This study |
| COL Δ*brxAB::*pRB473-*brx-roGFP2* | | This study |
| COL Δ*brxAB ypdA::*pRB473-*brx-roGFP2* | | This study |
| COL pRB473-*tpx-roGFP2* | | This study |
| COL Δ*ypdA::*pRB473-*tpx-roGFP2* | | This study |
| COL Δ*brxAB::*pRB473-*tpx-roGFP2* | | This study |
| COL Δ*ypdA::*pRB473-*ypdA* | | This study |
| COL Δ*brxAB::*pRB473-*brxA* | | This study |
| COL Δ*brxAB::*pRB473-*brxB* | | This study |
| *Staphylococcus* phage 81 | | [5] |

**Table S2. Plasmids**

| Plasmid | Description | Reference |
|---|---|---|
| pET11b | *E. coli* expression plasmid | Novagen |
| pET11b-*ypdA* | pET11b-derivative for overexpression of His-tagged YpdA | This work |
| pET11b-*ypdAC14A* | pET11b-derivative for overexpression of His-tagged YpdAC14A | This work |
| pET11b-*brxA* | pET11b-derivative for overexpression of His-tagged BrxA | [6] |
| pET11b-*gapDH* | pET11b-derivative for overexpression of His-tagged GapDH | [6] |
| pET11b-*brx-roGFP2* | pET11b-derivative for overexpression of His-tagged Brx-roGFP2 | [4] |
| pET11b-*tpx-roGFP2* | pET11b-derivative for overexpression of His-tagged Tpx-roGFP2 | This study |
| pRB473 | pRB373-derivative, *E. coli/ S. aureus* shuttle vector, containing xylose-inducible $P_{Xyl}$ promoter, $Amp^r$, $Cm^r$ $Amp^r$, $Cm^r$ | [7, 8] |
| pRB473-*brx-roGFP2* | pRB473-derivative expressing *brx-roGFP2* under $P_{Xyl}$ | [4] |
| pRB473-*tpx-roGFP2* | pRB473-derivative expressing *tpx-roGFP2* under $P_{Xyl}$ | This study |
| pRB473-*ypdA* | pRB473-derivative expressing *ypdA* under $P_{Xyl}$ | This study |
| pRB473-*brxA* | pRB473-derivative expressing *brxA* under $P_{Xyl}$ | This study |
| pRB473-*brxB* | pRB473-derivative expressing *brxB* under $P_{Xyl}$ | This study |

**Table S3. Oligonucleotide primers**

| Primer name | Sequence (5’ to 3’) |
|---|---|
| pET-tpx-for-NheI | CTAGCTAGCATGACTGAAATAACATTCAAAGG |
| pET-tpx-rev-SpeI | GCGACTAGTAATATTTTTGTATGCAGCTAAAGC |
| pET-ypdA-for-NdeI | GGAATTCCATATGCAAAAAGTTGAAAGTATCATA |
| pET-ypdAC14A-for-NdeI | GGAATTCCATATGCAAAAAGTTGAAAGTATCATAATTGGTGGAGGGCCA**GCG**GGATT |
| pET-ypdA-rev-BamHI | CGCGGATCCTTAGTGATGGTGATGGTGATGTGATTCTAAGGGCGTTTGTTTC |
| pRB-tpx-roGFP2-for-BamHI | CGCGGATCCTTAGTGATGGTGATGGTGATGTGATTCTAAGGGCGTTTGTTTC |
| pRB-tpx-roGFP2-rev-SacI | CGCGAGCTCTTACTTGTACAGCTCGTCCATGC |
| pMAD-ypdA-for-BglII | CGCAGATCTGACATACAGTGAATGGTCAAG |
| pMAD-ypdA-f1-rev | GTTATTTAGTACATAGACCTTTATTTCATTGTTTCGGCCTCCTTTAATC |
| pMAD-ypdA-f2-for | GATTAAAGGAGGCCGAAACAATGAAATAAAGGTCTATGTACTAAATAAC |
| pMAD-ypdA-rev-SalI | CCAGTCGACTGTATTGACAAAGGATCGTGTG |
| pMAD-brxA-for-BglII | CGCAGATCTCGATCATTTCGTGTATTTCCTA |
| pMAD-brxA-f1-rev | TAAATATGAATGCATATGATGCTCCTTTGACGAAAATTGTAAATAGT |
| pMAD-brxA-f2-for | ACTATTTACAATTTTCGTCAAAGGAGCATCATATGCATTCATATTTA |
| pMAD-brxA-rev-SalI | CCAGTCGACTGGAAGACTCGATTACGAATG |
| pMAD-brxB-for-BglII | CGCAGATCTCAATCGCAATGGTATCTTCATA |
| pMAD-brxB-f1-rev | GGATAGGTGATTGAACTTATGGATTGTGAAGAAAGATAAGAGGC |
| pMAD-brxB-f2-for | GCCTCTTATCTTTCTTCACAATCCATAAGTTCAATCACCTATCC |
| pMAD-brxB-rev-SalI | CCAGTCGACTGATATGATTGCAATTCGTAAC |
| pRB-ypdA-for-BamHI | TAGGGATCCTTAAAGGAGGCCGAAACAATGCAAAAAGTTGAAAGTATCA |
| pRB-ypdA-rev-KpnI | CTCGGTACCTTATGATTCTAAGGGCGTTTG |
| pRB-brxA-for-BamHI | TAGGGATCCTAATTGGAGGAATTAAATATGAAT |
| pRB-brxA-rev-KpnI | CTCGGTACCCTATTTACAATTTTCGTCAAAGG |
| pRB-brxB-for-BamHI | TAGGGATCCGGATAGGTGATTGAACTTATGG |
| pRB-brxB-rev-KpnI | CTCGGTACCTTATCTTTCTTCACAATATTTATTG |
| ypdA-NB-for | GGGCCATGCGGATTAAGTG |
| ypdA-NB-rev | CTAATACGACTCACTATAGGGAGACGTTCCCTGCAGCAATTACA |
| brxA-NB-for | GCTTATATGAAAGAAATTGCGC |
| brxA-NB-rev | CTAATACGACTCACTATAGGGAGAAATTTTCGTCAAAGGCATCCTT |
| brxB-NB-for | TTATACATGAACGGTGTTGTAG |
| brxB-NB-rev | CTAATACGACTCACTATAGGGAGAATTACGTTCATCACATCATGAC |

Restriction sites are underlined and bold bases indicate a point mutation.
