## Supplemental Figures S1-S6 for "*Staphylococcus aureus* uses the bacilliredoxin (BrxAB)/ bacillithiol disulfide reductase (YpdA) redox pathway to defend against oxidative stress under infections"

### Figure S1

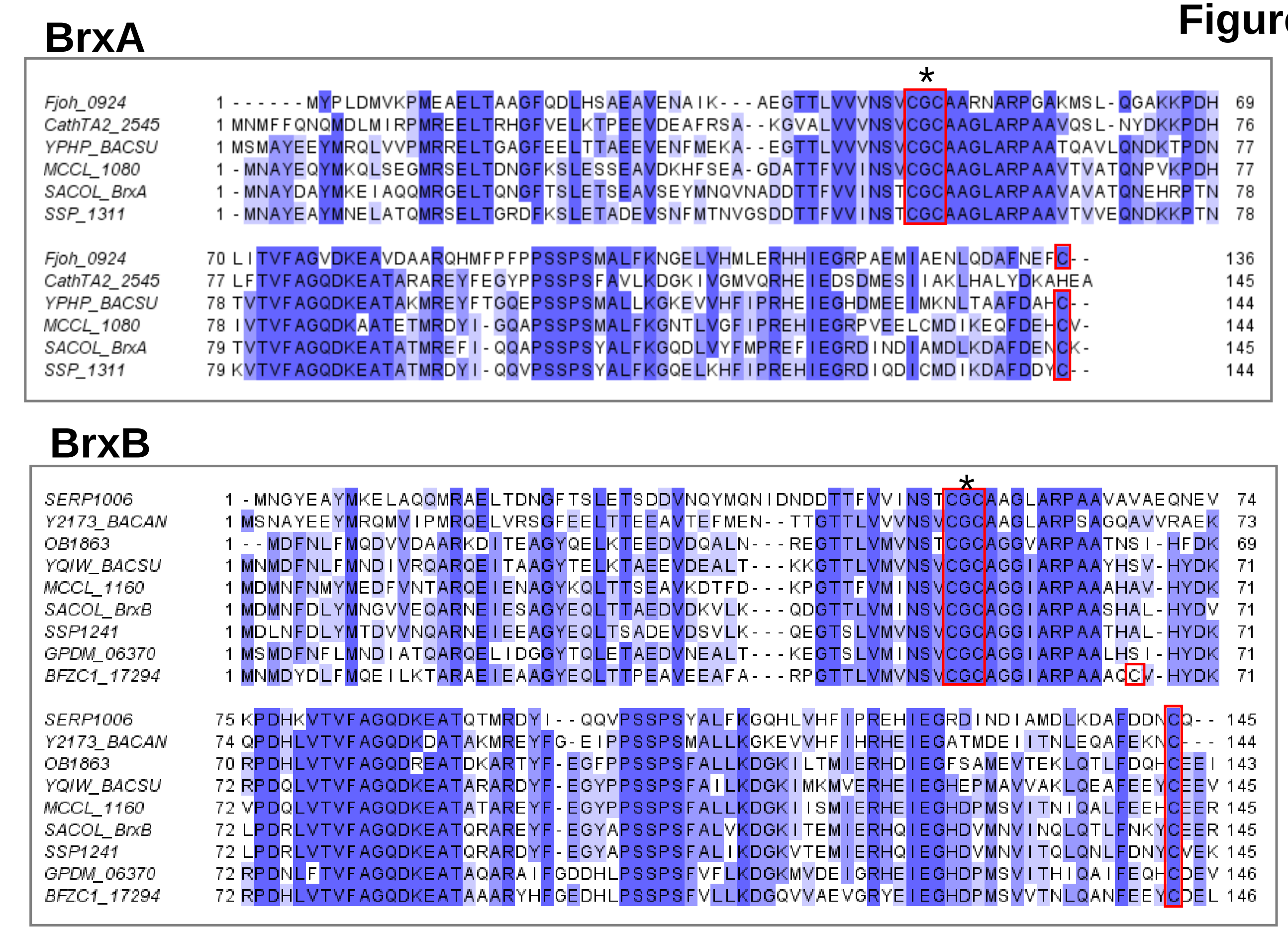


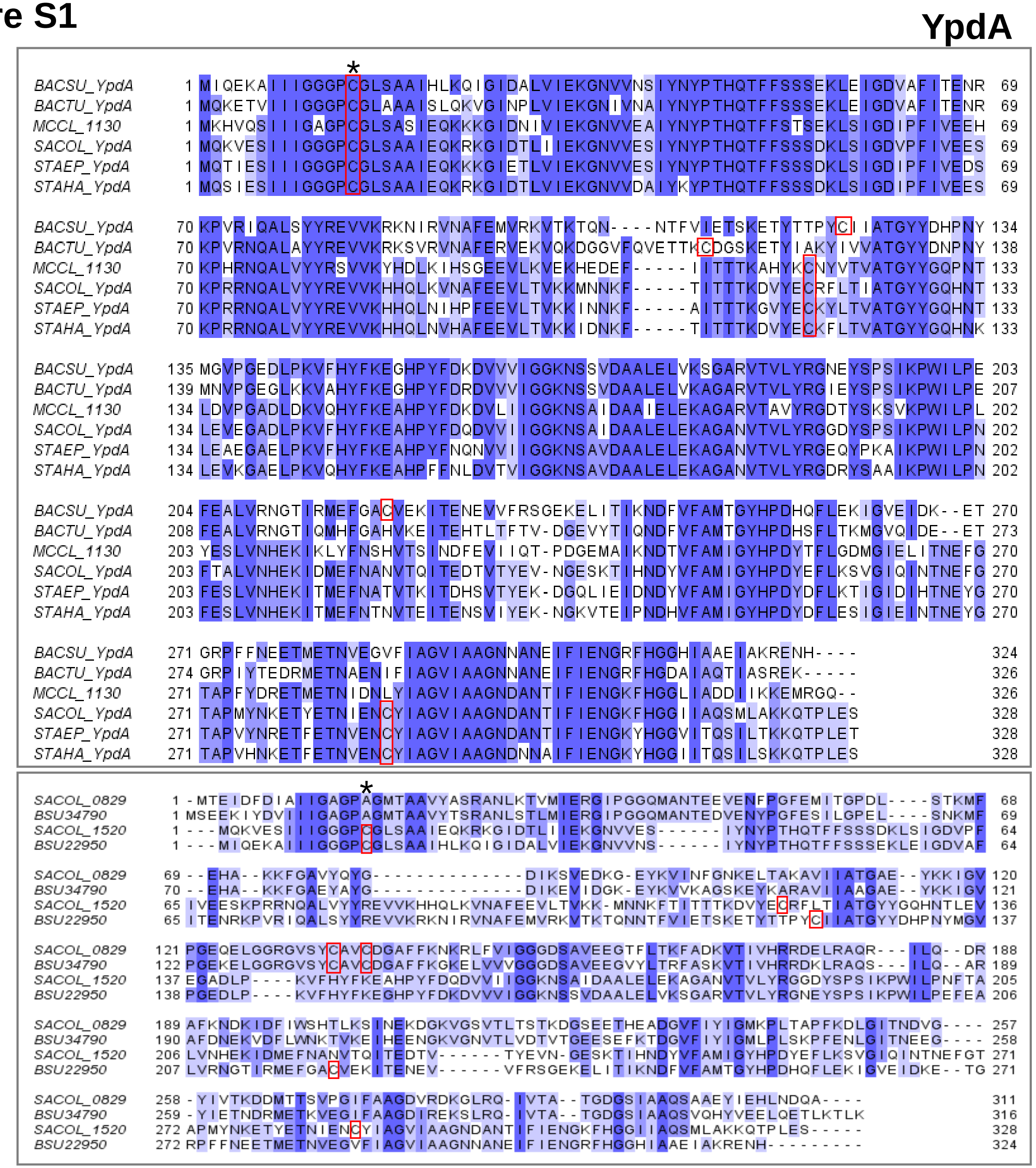


**Fig. S1. Multiple protein sequence alignments of BrxA/B and YpdA homologs across firmicutes.** The protein sequence alignment was performed with ClustalΩ2 and is presented in Jalview. The following protein sequences were aligned and the % identity to BrxA, BrxB and YpdA of *S. aureus* COL is given in parenthesis:

- **BrxA (SACOL1464)** of *S. aureus* COL, SSP1311 of *Staphylococcus saprophyticus* (73.1%), MCCL_1080 of *Macrococcus caseolyticus* (64.8%), BrxA (YPHP_BACSU) of *Bacillus subtilis* (55.1%), Fjoh_0924 of *Flavobacterium johnsoniae* (43.8%) and CathTA2_2545 of *Caldalkalibacillus thermarum* (42.6%).
- **BrxB (SACOL1558**) of *S. aureus* COL, SSP1241 of *S. saprophyticus* (84.1%), BrxB (YQIW_BACSU) of *B. subtilis* (68.3%), OB1863 of *Oceanobacillus iheyensis* (62.8%), BFZC1_17294 of *Lysinibacillus fusiformis* (59.6%), GPDM_06370 of *Planococcus donghaensis* (58.9%), Y2173_BACAN of *Bacillus anthracis* (46.3%) and SERP1006 of *Staphylococcus epidermidis* (43.3%).
- **YpdA (SACOL1520)** of *S. aureus* COL, YpdA (A0A2T4SSC4_STAHA) of *Staphylococcus haemolyticus* (82.6%), YpdA (A0A2G7I444_STAEP) of *S. epidermidis* (82.3%), YpdA (MCCL_1130) of *M. caseolyticus* (68.6%), YpdA (YPDA_BACSU) of *B. subtilis* (62.8%) and YpdA (A0A0Q0R6S1_BACTU) of *Bacillus thuringiensis (*60.1%). In the lower panel, YpdA of *S. aureus* (SACOL1520) and *B. subtilis* (BSU22950) were aligned with TrxB of *S. aureus* (SACOL0829) and TrxB of *B. subtilis* (BSU34790). The conserved CGC motif of BrxA/B and the conserved Cys14 residue of YpdA are labeled in red with an asterisk (*).

#### Figure S2

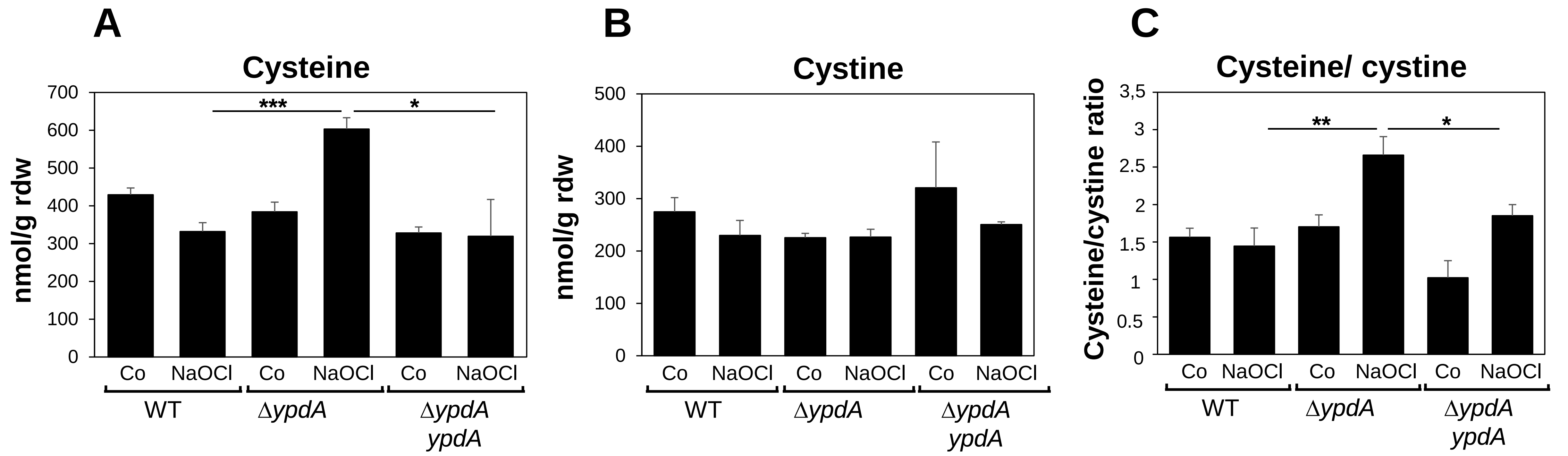


**Figure S2. Determination of the cysteine and cystine levels before and after NaOCl stress in the *S. aureus* COL wild type (WT), the *ypdA* mutant and *ypdA* complemented strain.** *S. aureus* COL strains were grown in RPMI medium until an $OD_{500}$ of 0.9 and exposed to 2 mM NaOCl stress for 30 min. The cysteine and cystine levels were determined after mBBr derivatisation of LMW thiols and disulfides and measured by HPLC thiol metabolomics. Mean values are presented with standard deviation of three biological replicates. $^{ns}p > 0.05$; $^{*}p \leq 0.05$ $^{**}p \leq 0.01$ and $^{***}p \leq 0.001$.

**Figure S3**

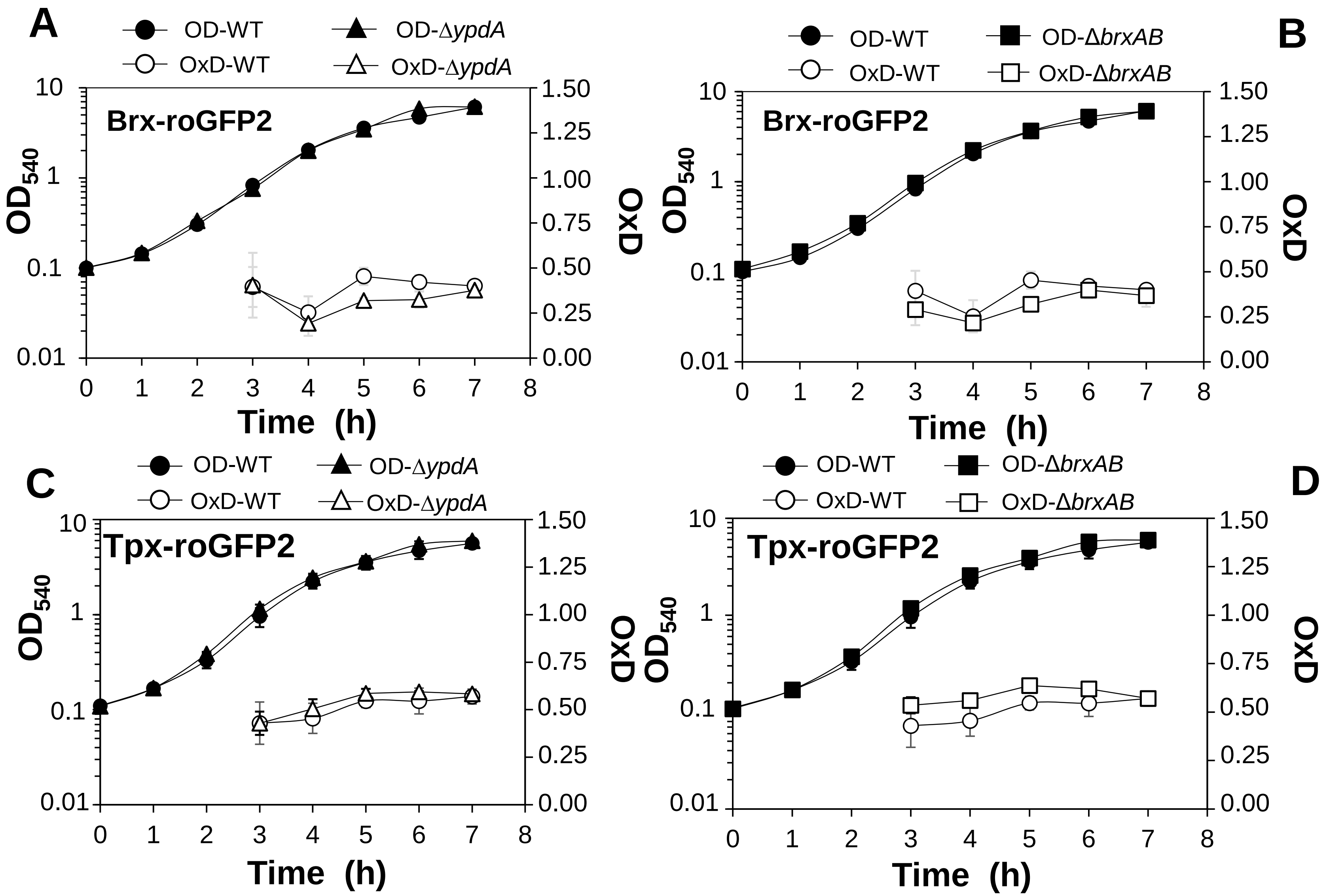


**Fig. S3. The basal level of the BSH redox potential is not affected in *ypdA* mutant during the growth as determined with the Brx-roGFP2 biosensor. (A,B)** The basal level oxidation of Brx-roGFP2 is similar in the *S. aureus* COL WT, *ypdA* and *brxAB* mutants along the growth curves in LB medium, indicating that the Brx/YpdA redox pathway is not essential to maintain the BSH redox potential during the growth. **(C,D)** Oxidation of the Tpx-roGFP2 biosensor is similar in the *S. aureus* COL WT, *ypdA* and *brxAB* mutants along the growth curves in LB medium. This indicates no increased levels of endogenous ROS formation in the absence of the Brx/BSH/YpdA pathway. In all graphs, mean values are presented and error bars indicate the standard deviation of three biological replicates.

**Figure S4**

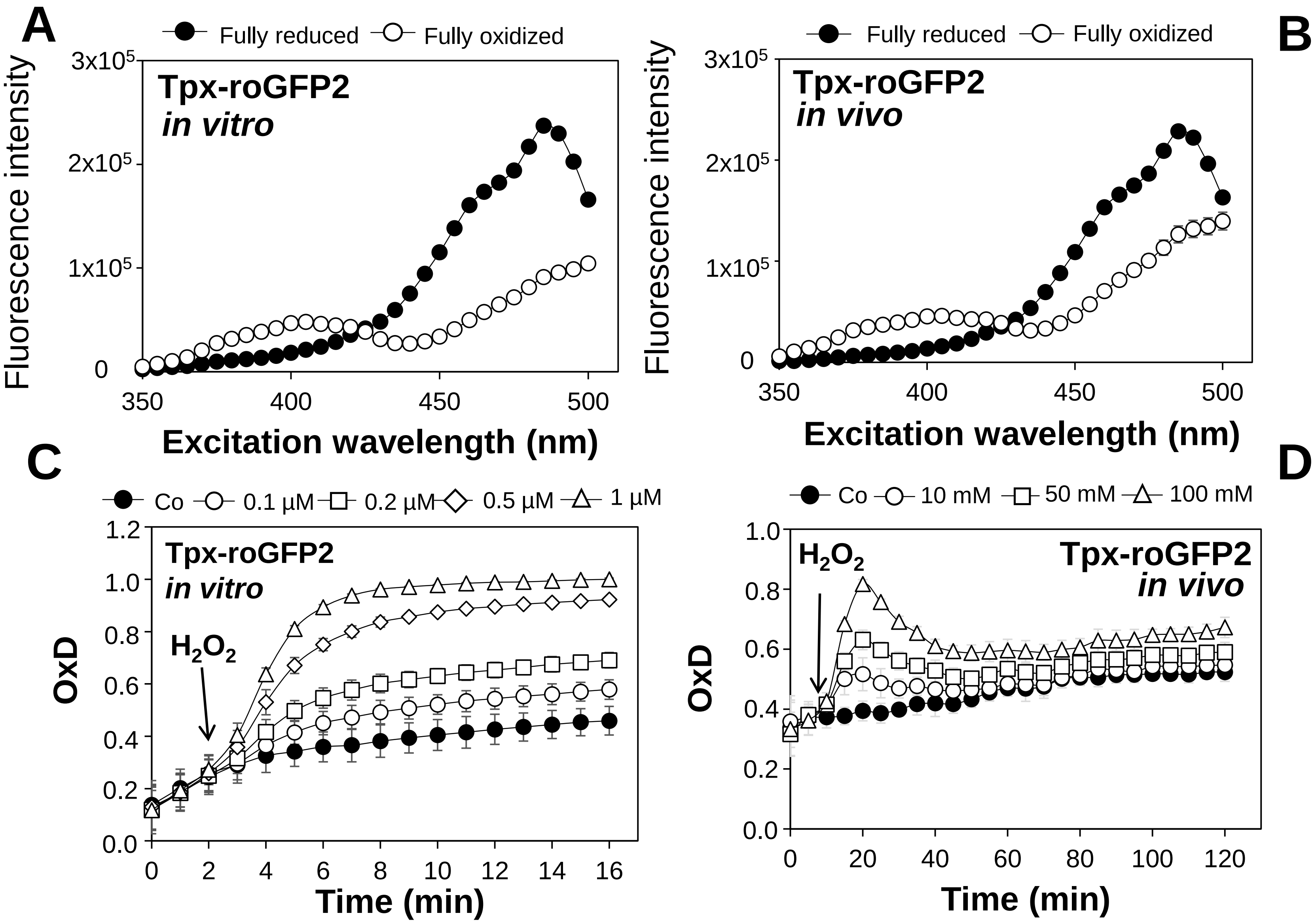


**Fig. S4. Responses of Tpx-roGFP2 *in vitro* (A,C) and *in vivo* inside *S. aureus* COL after exposure to $H_2O_2$. (B, D)** The ratiometric Tpx-roGFP2 biosensor response in the DTT-treated fully reduced and diamide-treated fully oxidized state *in vitro* **(A)** and inside *S. aureus* COL *in vivo* **(B)**. For fully reduced and oxidized Tpx-roGFP2, 10 mM DTT and 5 mM diamide were used *in vitro* as well as 15 mM DTT and 20 mM cumene hydroperoxide (CHP) *in vivo* (n = 5). **(C)** The purified Tpx-roGFP2 biosensor (1 µM) responds specifically to low levels $H_2O_2$ (0.1-1 µM $H_2O_2$) *in vitro*. **(D)** Tpx-roGFP2 inside *S. aureus* COL is rapidly and reversible oxidized by sub-lethal 1-100 mM $H_2O_2$ *in vivo*. The OxD was calculated based on the 405/488 nm excitation ratio with emission measured at 510 nm. Mean values with standard deviations are shown in all graphs.

**Figure S5**

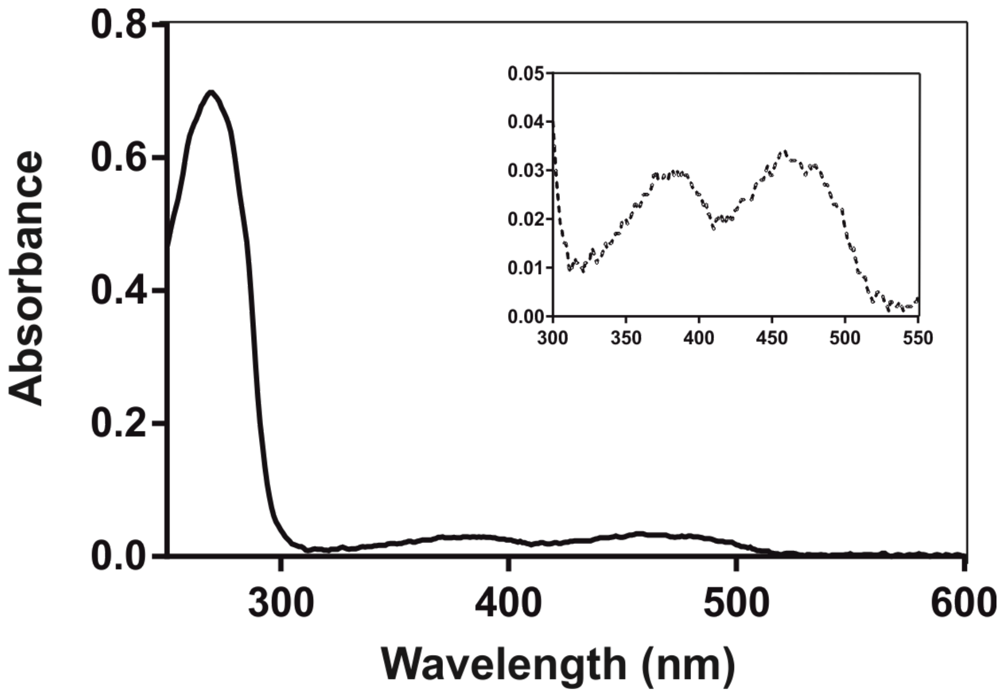


**Fig. S5.** The UV-visible absorption spectrum of purified yellow coloured YpdA protein indicates that YpdA is a flavoprotein containing the FAD co-factor The insert indicates the FAD cofactor of YpdA with absorbance peaks at 375 and 450 nm in the UV-visible spectrum.

**Figure S6**

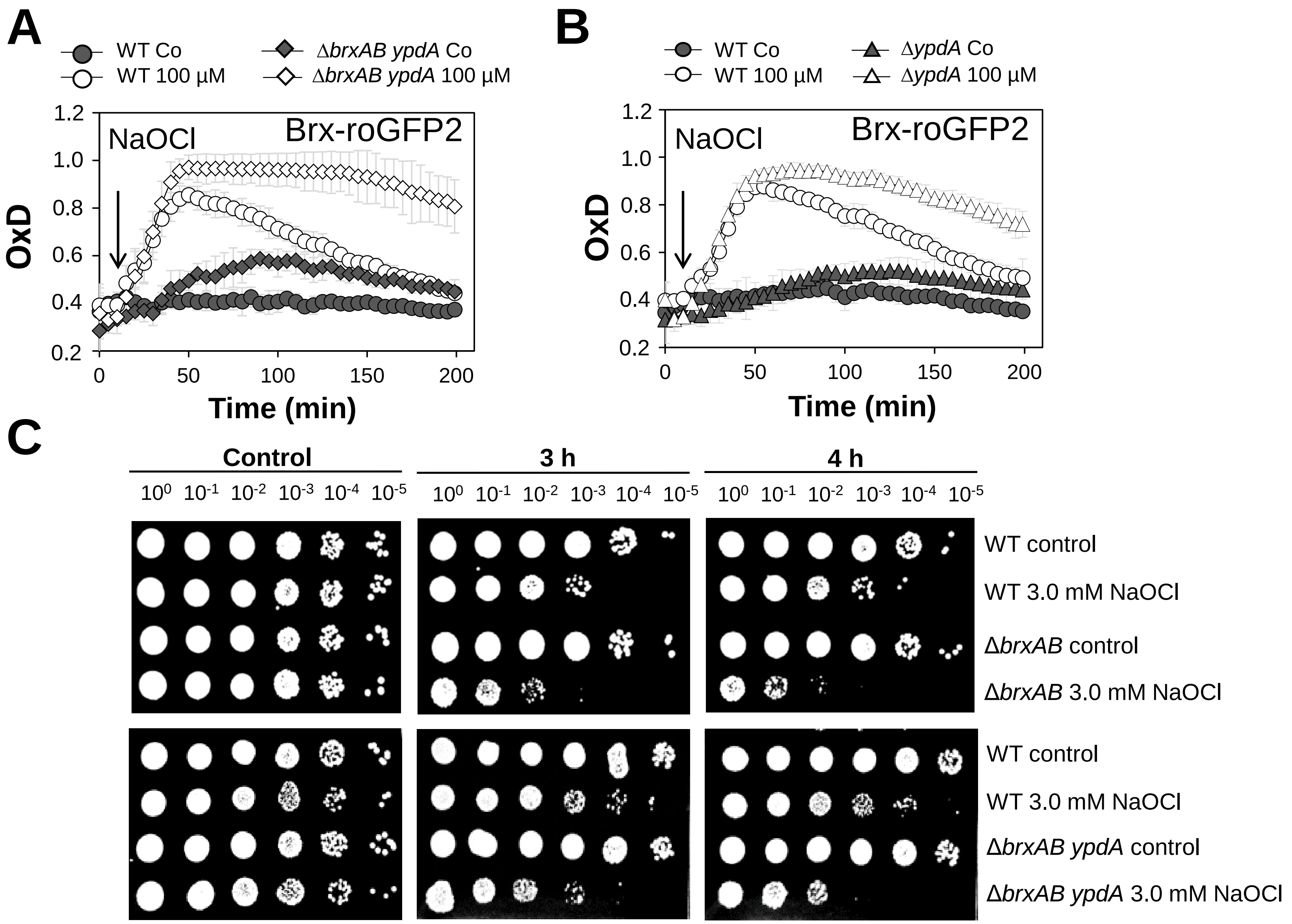


**Fig. S6. Brx-roGFP2 measurements and survival assays indicate that the *S. aureus brxAB ypdA* triple mutant is similar defective to rescue $E_{BSH}$ as the *ypdA* mutant and similar sensitive to NaOCl stress as the *brxAB* mutant. (A, B)** Brx-roGFP2 response in the *S. aureus* COL WT, *ypdA* and *brxAB ypdA* mutants under 100 µM NaOCl stress**.** Both mutants show the same defects to recover reduced $E_{BSH}$. **(C)** Qualitative survival assays of the *S. aureus* COL WT, *brxAB* and *brxAB ypdA* mutants under 3.0 mM NaOCl stress. Survival was analysed by spotting 10 µl of serial dilutions of *S. aureus* cells after different hours of NaOCl-exposure on agar plates, which were incubated over night at 37°C. The results indicate that the *brxAB ypdA* triple mutant displays the same sensitivity as the *brxAB* double mutant.
